# The role of geology in creating stream climate-change refugia along climate gradients

**DOI:** 10.1101/2022.05.02.490355

**Authors:** Nobuo Ishiyama, Masanao Sueyoshi, García Molinos Jorge, Kenta Iwasaki, N Junjiro Negishi, Itsuro Koizumi, Shigeya Nagayama, Akiko Nagasaka, Yu Nagasaka, Futoshi Nakamura

## Abstract

Identifying climate-change refugia is a key adaptation strategy for reducing global warming impacts. Knowledge of the effects of underlying geology on thermal regime along climate gradients and the ecological responses to the geology-controlled thermal regime is essential to plan appropriate climate adaptation strategies. The dominance of volcanic rocks in the watershed is used as a landscape-scale surrogate for cold groundwater inputs to clarify the importance of underlying geology. Using statistical models, we explored the relationship between watershed geology and the mean summer water temperature of mountain streams along climate gradients in the Japanese archipelago. Summer water temperature was explained by the interaction between the watershed geology and climate in addition to independent effects. The cooling effect associated with volcanic rocks was more pronounced in streams with less summer precipitation or lower air temperatures. We also examined the function of volcanic streams as cold refugia under contemporary and future climatic conditions. Community composition analyses revealed that volcanic streams hosted distinct stream communities composed of more cold-water species compared with non-volcanic streams. Scenario analyses revealed a geology-related pattern of thermal habitat loss for cold-water species. Non-volcanic streams rapidly declined in thermally suitable habitats for lotic sculpins even under the lowest emission scenario (RCP 2.6). In contrast, most volcanic streams will be sustained below the thermal threshold, especially for low and mid-level emission scenarios (RCP 2.6, 4.5). However, the distinct stream community in volcanic streams and geology-dependent habitat loss for lotic sculpins was not uniform and was more pronounced in areas with less summer precipitation or lower air temperatures. Although further studies are needed to understand underlying mechanisms of the interplay of watershed geology and climate, findings highlight that watershed geology, climate variability, and their interaction should be considered simultaneously for effective management of climate-change refugia in mountain streams.

## 1. Introduction

Global warming represents an extreme threat to the world’s biodiversity (Thomas et al. 2004). The global surface temperature has already increased by 1.09 °C on average since the pre-industrial era (IPCC 2021) and the current cumulative greenhouse gas emissions continue to track the most aggressive projection scenarios (Schwalm et al. 2020). One adaptation strategy for minimizing the dire impacts of climate change, including global warming, on ecosystems is to identify and manage climate-change refugia (Morelli et al. 2016, Morelli et al. 2020). Climate-change refugia are areas that mitigate the larger scale impacts of climate change over time and enable the persistence of valued physical and ecological resources (Morelli et al. 2016). In the context of global warming, understanding the driving factors for local thermal heterogeneity in landscapes is the first step for the effective management of climate-change refugia.

There are multiple surface and subsurface factors that create thermal heterogeneity in landscapes. Factors encompassing topography such as elevation, slope aspect, and topographic convergence are key drivers of terrestrial or aquatic thermal heterogeneity in mountain regions (Dobrowski 2011, Isaak et al. 2016, Macek et al. 2019, Ackerly et al. 2020). The cooling effects associated with forest canopy cover is also widely reported in the terrestrial and freshwater realms (Nagasaka and Nakamura 1999, Haesen et al. 2021). Extensive monitoring networks in European forests revealed that subcanopy air temperatures were cooler than the free air temperatures by 2.0 °C on average (Haesen et al. 2021). The Earth’s subsurface factors, namely the geology, also contribute to local thermal heterogeneity. However, studies are limited compared to those examining the role of the Earth’s surface factors. In freshwater, the watershed geology controls the thermal heterogeneity by providing inputs of cold groundwater (e.g., Tague et al. 2007, McDonnell et al. 2015, Lusardi et al. 2021). In the context of Antarctic active layers, Hrbáček et al. (2017) reported a distinct ground thermal regime between lithologically different sites due to thermal conductivity controlled by the geology. More importantly, effect of these covariates on local thermal conditions are not always uniform and can vary along environmental gradients (Hilderbrand et al. 2014, Jucker et al. 2018, Ehbrecht et al. 2019, De Frenne et al. 2021). These potential interactions among landscape and climatic factors imply that identification and management of climate-change refugia based on independent effects cannot effectively mitigate biodiversity decline in the future. However, prior knowledge on interactive effects shaping thermal heterogeneity in landscapes has placed a disproportionate emphasis on the Earth’s surface factors such as topography and land cover, with less focus on subsurface factors. Given that global warming and biodiversity decline have occurred over various underlying geologies, there has been a growing interest in conserving nature’s stage in the context of environmental heterogeneity for biodiversity conservation (e.g., Knudson et al. 2018, Gray 2021). Knowledge of the effects of underlying geology on the thermal regime along environmental gradients is essential to deepening our understanding of the role of nature’s stage in climate change adaptation.

River water temperature is globally increasing as well as atmospheric temperature (Webb and Nobilis 2007, Kaushal et al. 2010). Evidence is growing for distribution shifts and community reorganization due to the warming trend in running water ecosystems (Chessman 2009, Comte and Grenouillet 2013, Comte et al. 2021). High spatiotemporal thermal heterogeneity characterizes running water (Caissie 2006, Tonolla et al. 2010). Groundwater inputs with a stable thermal regime are one of the key factors shaping thermal refugia at multiple spatial scales (Kanno et al. 2014, Snyder et al. 2015, Hare et al. 2021). Watershed geology affecting water permeability in the subsurface layers contributes to groundwater inputs (Mushiake et al. 1981, Boulton and Hancock 2006, Cornu et al. 2013). Thermal heterogeneity controlled by geology in running water ecosystems has been increasingly highlighted (Tague et al. 2007, McDonnell et al. 2015, Lusardi et al. 2021). In addition to the cooling process at the streambed/water interface, namely the groundwater contribution, climatic conditions, such as air temperature and precipitation, determine the water temperature (Caissie 2006). However, previous studies have not clearly shown the importance of watershed geology along climate gradients. Identification and management of climate-change refugia is especially important in river networks, where the movement of aquatic organisms such as fish and larval insects is highly confined to network branches (Campbell Grant et al. 2007). Considering the increased exposure and impaired capacity of aquatic organisms to respond to the effects of climate warming, testing the geology–climate interaction has the potential to advance climate change adaptation measures in running water ecosystems.

In the present study, we explored the relationship between the watershed geology and the summer water temperature of mountain streams along climate gradients in the Japanese archipelago, including air temperature and precipitation. Japan provides an ideal study area for this purpose due to its diverse geology and climate. Two oceanic plates subduct below two continental plates around the Japanese archipelago (Sato 1994), creating a complex geological landscape with a variety of lithologies. Japanese geology is also characterized by continuous volcanism. The subduction boundaries activate magma derived from the mantle, resulting in active volcanos. The high spatiotemporal variability in the Japanese archipelago of temperature and precipitation is associated with the difference in geographic features extending from south to north, complex topography, and varying ocean currents. In this study, we focused on the dominance of volcanic rocks in the watershed as a landscape-scale surrogate for groundwater inputs. This is because they support groundwater discharge more than other bedrock types due to their high permeability (Tague and Grant 2004, Fujimoto et al. 2016, Iwasaki et al. 2021).

Given the continuous impacts of global warming, streams providing cold habitats will act as climate-change refugia for cold-water species (Isaak et al. 2016). However, the importance of watershed geology in creating climate-change refugia and its climate dependency have received little attention in previous studies. In the present study, we further examined the function of volcanic watersheds as cold refugia under contemporary and future climatic conditions in areas with different climate conditions. Various species use cold water habitats created by groundwater inputs as thermal refugia (Brewitt and Danner 2014, Kanno et al. 2014) and contribute to sustaining source populations (Nakajima et al. 2021). Therefore, we hypothesize that stream communities differ between non-volcanic and volcanic streams, and that the difference is attributed to more cold-water species in volcanic streams. We also predicted suitable thermal habitats for cold water species under future climatic conditions to examine the geology-related patterns of loss of suitable thermal habitats. We postulated that streams with a high level of volcanic rock would function as climate-change refugia for cold-water species due to the high cooling effect of groundwater inputs. To clarify the importance of watershed geology in creating cold refugia and their climate dependency, these hypotheses were tested in regions with contrasting climate conditions. The study area encompassed northern Japan which has low summer air temperatures and low precipitation and central Japan that has high summer air temperatures and high precipitation.

## 2. Materials and Methods

### 2.1. Modeling of stream summer water temperature

#### 2.1.1 Study Sites

Five study regions were selected from middle to high latitudes in Japan to construct a temperature monitoring network encompassing a range of climatic conditions (Fig. 1, Table 1). We set paired air-water temperature loggers in the mountain streams in the five study regions to model the mean summer water temperature. The monitoring sites were between 35.4° and 43.8° latitude, 136.4° and 143.8° longitude (WGS84), with an altitude ranging from 28–1233 m. We selected low order (area of watershed < 100 km^2^) and low disturbance (forest cover ratio in the watershed of > 80%) streams as study sites. All the streams belong to independent rather than nested watersheds to avoid the cooling or warming effect of upstream sites on downstream sites. The mean drainage area, site elevation, and percentage of forest cover within the watersheds were 9.5 ± 10.9 km^2^, 425.6 ± 240.2 m, and 96.6 ± 4.3% (mean ±SD), respectively. The watershed geology of the streams in this study included volcanic, sedimentary, metamorphic, plutonic rocks, and accretionary complexes (Fig. 1). The volcanic rocks in the watersheds are mainly non-alkaline mafic volcanic, pyroclastic flow volcanic, and felsic volcanic rocks, with low proportions of volcanic debris and non-alkaline felsic volcanic intrusive rocks (Fig. S1). The mean proportion of the volcanic rocks in the watershed, summer air temperature, and the total summer precipitation was 43.7 ± 45.0%, 19.8 ± 1.9 °C, and 476.5 ± 362.9 mm (mean ± SD), respectively.

**Fig. 1.**
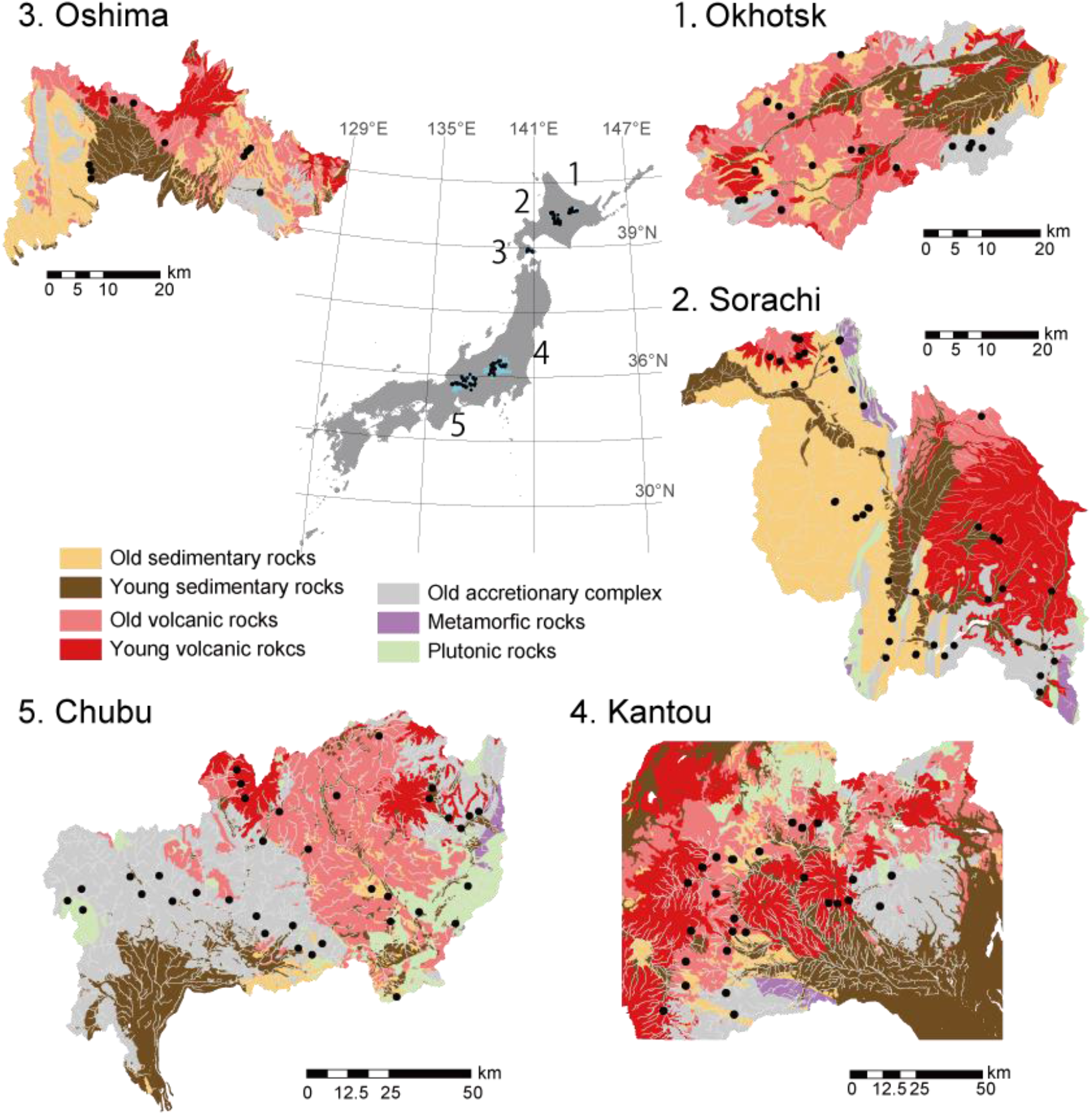
Water and air temperature monitoring streams in the middle to high latitudes of Japan. The number of study streams for the water temperature modeling is 22, 44, 11, 27, and 30 in Okhotsk, Sorachi, Oshima, Kantou, and Chubu, respectively. In the geological legend, “old” means pre-Quaternary, and “young” means Quaternary. The geological map is created based on the lithology classification of the Seamless Digital Geological Map of Japan (Geological Survey of Japan).

**Fig. 2.**
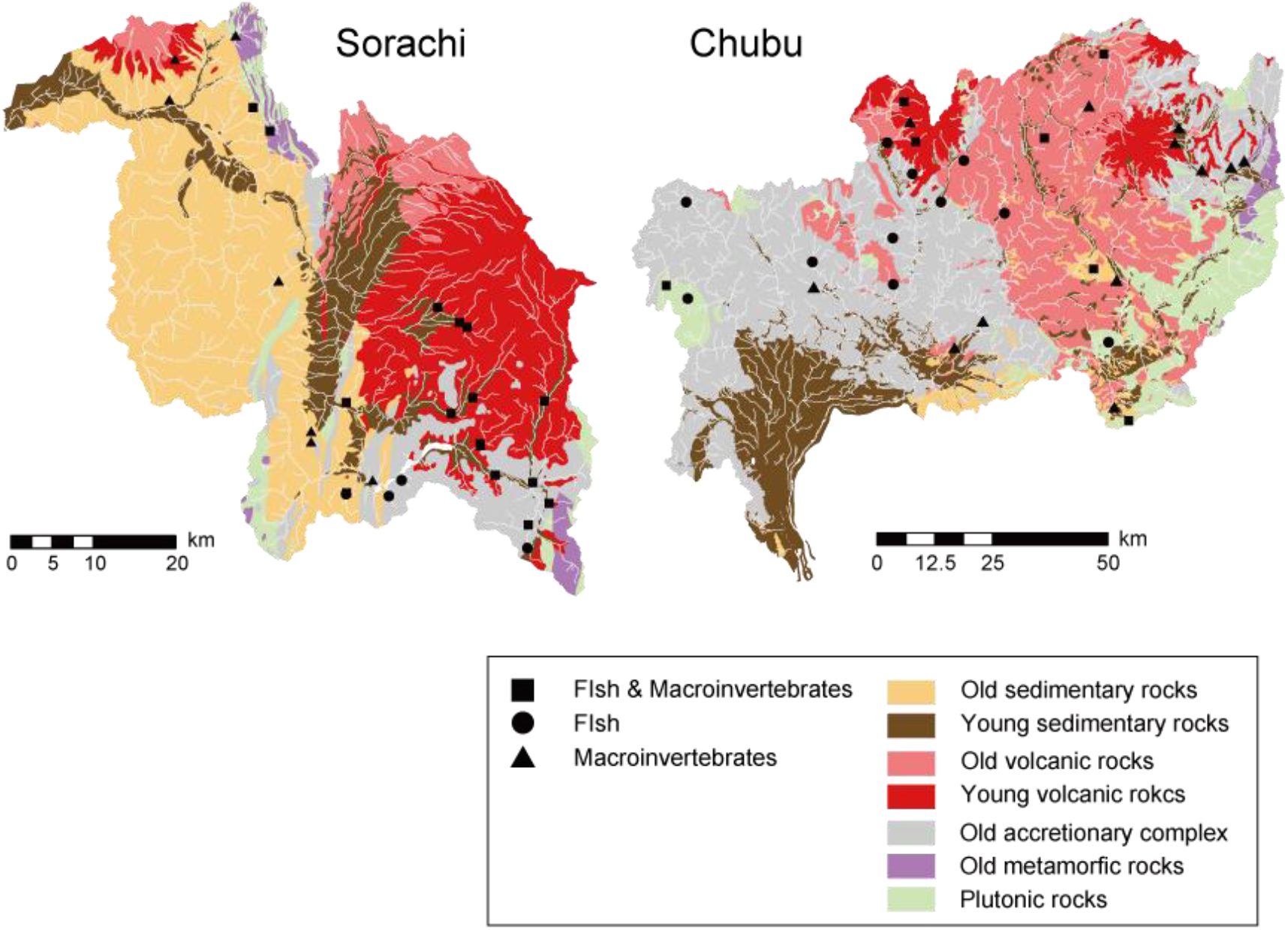
Location of the biological monitoring streams in a) the Sorachi region and b) the Chubu region.

**Table 1.** Characteristics of the five study regions. The average values are shown for the monitoring sites.

| Region | Total summer precipitation (mm) | Mean summer air temperature (°C) | Agricultural land use in watershed (%) | Elevation (m) | Drainage area (km<sup>2</sup>) | Volcanic in watershed (%) |
| --- | --- | --- | --- | --- | --- | --- |
| Okhotsk | 151.7 | 18.1 | 0.0 | 453.3 | 6.7 | 51.0 |
| Sorachi | 280.0 | 18.6 | 0.3 | 333.4 | 8.6 | 32.8 |
| Oshima | 334.4 | 20.3 | 0.0 | 110.5 | 4.3 | 55.2 |
| Kantou | 440.7 | 21.1 | 0.8 | 614.3 | 8.9 | 63.5 |
| Chubu | 990.8 | 21.4 | 0.9 | 483.3 | 13.3 | 29.3 |

#### 2.1.2 Monitoring of air and water temperatures

Air and water temperatures at each site were recorded hourly using the Onset Hobo UA-001-64 (Hobo, Lakeville, MA, USA) (accuracy ± 0.53 °C), Elitech RC-5 (± 0.5 °C) (Elitech, Paris, France), or Gemini TG-4100 (± 0.5 °C) (Gemini, New York, NY, USA) temperature data loggers. Water loggers were installed in the non-turbulent flowing sections of the stream housed in short sections of perforated PVC pipes to facilitate water exchange and to shield them from sunlight. Air loggers on perforated PVC pipes were attached to nearby trees (approximately 2 m above ground) shaded by canopy covers in riparian areas. We calculated the summer mean air and water temperature using thermal data from July and August. Sites with a high proportion of missing values (> 10 days in the summer season) were excluded from the analysis due to problems such as equipment failures, air exposure, and loss of loggers associated with floods. A total of 290 observations obtained from 140 stream sites were used for analysis with a mean monitoring duration of 2.1 summer seasons per site.

#### 2.1.3 Covariates for water temperature modeling

Seven covariates that could affect the mean summer water temperature were applied, namely the proportion of volcanic rocks and agricultural land in the watershed, the drainage area, the stream slope, the site elevation, the mean summer air temperature, and the total summer precipitation. The pairwise interactions of the volcanic rocks in the watershed with the mean summer air temperature and the total summer precipitation were used to test the importance of watershed geology along climate gradients. The watershed geology and land use were calculated based on the 1:200,000 Seamless Digital Geological Map of Japan provided by the Geological Survey of Japan (https://gbank.gsj.jp/seamless/2d3d/) and the ALOS/AVNIR-2 High-Resolution Land Cover map (2018-2020) provided by the Japan Aerospace Exploration Agency (https://www.eorc.jaxa.jp/ALOS/jp/dataset/lulc/lulc_v2111_j.htm), respectively. The drainage area (km^2^), stream slope, and site elevation were obtained from a 10 m digital elevation model published by the Geospatial Information Authority of Japan. For calculating the stream slope, we targeted a 200 m stream reach centered on a study site. For the total summer precipitation, we collected daily precipitation data at each study monitoring site from the 1 km spatial resolution Agro-Meteorological Grid Square Data, NARO (Ohno et al. 2016) and calculated the total summer precipitation. All the geospatial analyses were performed using ArcGIS Pro (Esri, version 2.4.0).

#### 2.1.4 Statistical analyses

We constructed generalized linear mixed models with a Gaussian error distribution and an identity link function. The nested random intercept (site within the watershed) was applied for random effect. The response variable was the mean summer water temperature, and the covariates were above seven variables and two interactions (geology-air temperature and geology-precipitation). The proportion of agricultural land in the watershed was natural log-transformed to reduce the influence of a few outliers with high values. All the variables were used for the analyses because there was no high correlation between them (Fig. S2) and variance influence factors ranging from 1.1 to 2.4 showed a low risk of multicollinearity (VIF < 10, Neter et al. 1996). The model performance was evaluated using the Akaike information criterion corrected for the small sample size (AICc; Burnham and Anderson 2002). We conducted model averaging for superior models (ΔAICc < 2) using the Akaike weight of each model and included the final model. The parameters for which the 95% confidence intervals did not include zero were chosen as the influential factors. Goodness of fit for the final model was assessed using the conditional and marginal coefficients of determination R^2^ (Nakagawa and Schielzeth 2013, Nakagawa et al. 2017), representing the total variance explained by the model with fixed and random components together and the variance explained by the fixed effects. The root mean square error (RMSE) was also computed as a measure of the goodness-of-fit. We used R software version 4.1.1 (R Development Core Team 2021) for water temperature modeling with the glmmTMB (Brooks et al. 2017) and MuMIn packages (Barton 2022).

### 2.2. Responses of stream community composition to watershed geology

#### 2.2.1. Biotic survey

To test the hypothesis that streams with a higher proportion of volcanic rocks within the contributing watershed host stream communities characterized by more cold-water species, we collected stream fish and macroinvertebrates from the Sorachi and Chubu regions. We selected two regions with contrasting climate conditions to clarify the climate dependency of the contribution of watershed geology in determining stream communities (Table 1). In the Sorachi region, 26 riffles in 20 streams and 43 riffles in 22 streams were selected as sampling sites for fish and macroinvertebrates, respectively. Fish were sampled in July and August 2018, and macroinvertebrates was sampled in May and June 2021. In the Chubu region, we set 32 reaches in 18 streams and 19 riffles in 19 streams as sampling sites for fish and macroinvertebrates, respectively. Fish were collected between September to November from 2015 to 2017, and macroinvertebrates were collected from November 2020 to February 2021.We selected the streams used for biomonitoring based on accessibility, naturalness without a high correlation between watershed geology and other landscape-scale factors (Fig. S3). The fish were collected with two pass electrofishing using a backpack electrofisher (Model LR-20B; Smith-Root Inc., Vancouver, Washington, USA) to quantify the relative abundance of each species according to the species density (individuals/100 m^2^). Fish sampling was authorized by the Fisheries Adjustment Rules of the Hokkaido, Gifu and Nagano Prefectures. For the macroinvertebrate surveys, three quadrat samples were collected from each study riffle using a Surber net (0.0625 m^2^, mesh size 0.475 mm). Ephemeroptera, Plecoptera, Trichoptera, Diptera and Coleoptera were identified at the species level based on Kawai and Tanida (2005), except for certain taxa which were identified at the subfamily and family levels because of a lack of detailed taxonomic information. Three samples were pooled together and the total number of individuals for each species was recorded for each study riffle (individuals/0.1875 m^2^).

#### 2.2.2. Local environmental survey

The summer water temperature, water depth, current velocity, substrate coarseness, and water quality indices were measured to examine the influence of local environments on the stream community composition (Table S1). The mean summer water temperature was calculated for each study stream (see “Monitoring of air and water temperatures” for details). The local environments for fish and macroinvertebrates were measured separately to represent habitat conditions for each taxon. For the fish habitat, we established five equally spaced transects with five measurements for each study site. The water depth and current velocity were determined at each measurement point at a 60% depth, and the means of these variables for each riffle were used for the following analysis. Current velocity was measured using an electromagnetic current meter. At each measurement point, substrates were identified through visual observation and scored as one of the following types: bedrock (0), sand (1: < 2 mm), gravel (2: 2–16 mm), pebbles (3: 17–64 mm), cobbles (4: 64–256 mm), or boulders (5: > 256 mm). The recorded scores were averaged to represent the site scale substrate coarseness. Dissolved water oxygen (mg/L) was also measured for each study stream using water quality meters. For macroinvertebrate habitat, we measured the water depth and the current velocity at the four corners and at the center of the three quadrats and calculated the mean water depth and the current velocity. The substrate score was visually recorded at the same points to calculate the substrate coarseness. Surface water samples were collected from each stream studied and taken back to the laboratory in a cooler box with ice packs. NO3^-^ (mg/L) of each sample was analyzed using an ion chromatography system, Dionex ICS 1100 (DIONEX, USA). The methods used for the water sample processing were in line with those used by Iwasaki et al. (2021).

#### 2.2.3. Stream community composition and indicator species

Two-dimensional non-metric multi-dimensional scaling (NMDS) ordinations were conducted to examine the variability of fish and macroinvertebrate community composition between the geological types (volcanic vs. non-volcanic). Each study riffle or reach was applied as an analysis unit. The Bray–Curtis metric was applied to measure the dissimilarity of community composition based on log-transformed abundance data. NMDS was performed using the metaMDS function of the vegan package (Oksanen 2020). The fit of the reproduced similarity matrix to the observed similarity matrix was assessed using the stress value. Correlations between the local environments measured and the watershed geology to the ordination scores were evaluated using the envfit function in the Vegan package. Significant differences in the community composition between geological types were examined using permutational multivariate analysis of variance (PERMANOVA; 999 permutations) and the adonis function in the Vegan package. For the PERMANOVA, the stream type (volcanic or non-volcanic) was classified based on whether volcanic rocks represented the dominant watershed geology in the contributing catchment, accounting for over 50% of the catchment area.

We hypothesized that stream communities significantly differ between non-volcanic and volcanic streams due to abundant cold-water species in the volcanic streams. To test this hypothesis, we then conducted an indicator value analysis based on the abundance-based phi coefficient index of association as a fidelity value (De Cáceres and Legendre 2009) when we found a significant difference in the community composition between geological types. This analysis identifies species that are associated with the geological types. The multipatt function in the indicspecies package was used for the analyses (De Caceres et al. 2022). Based on the results of the indicator species analysis, thermal preference of the indicator species selected was evaluated using the wider literature (Please see the caption of Table 3 for details).

**Table 3.** Summary of the abundance-based indicator species analysis in the Sorachi region. Species with significant fidelity values are listed. The value of the correlation and the statistical significance of the association is shown as r_pb_ and the p-value, respectively. In the thermal tolerance classification, cold-water species and cool-warm water species are shown as 1 and 2, respectively. Cold-water species are indicated by bold letters. For macroinvertebrates, the top ten indicator species with a higher r_pb_ are shown.

| Taxa | $r_{pb}$ | p | Thermal tolerance <sup>a)</sup> | Taxa | $r_{pb}$ | p | Thermal tolerance <sup>a)</sup> |
| --- | --- | --- | --- | --- | --- | --- | --- |
| Non-volcanic streams |  |  |  | Volcanic streams |  |  |  |
| (Fish) |  |  |  | (Fish) |  |  |  |
| *Barbatula barbatula* | 0.444 | 0.028 * | 2 | ***Cottus nozawae*** | **0.512** | **0.01 \*\*** | **1** |
| (Macroinvertebrates) |  |  |  | (Macroinvertebrates) |  |  |  |
| *Hexatoma* spp. | 0.629 | 0.002 ** | 2 | **Perlodidae spp.** | **0.860** | **0.001 \*\*\*** | **1** |
| ***Rhithrogena japonica*** | **0.400** | **0.005 \*\*** | **1** | **Diamesinae spp.** | **0.699** | **0.001 \*\*\*** | **1** |
| Chloroperlidae spp. | 0.391 | 0.019 * | 2 | ***Brachycentrus americanus*** | **0.675** | **0.001 \*\*\*** | **1** |
| *Hydropsyche orientalis* | 0.340 | 0.034 * | 2 | ***Heterolimnius hasegawai*** | **0.629** | **0.001 \*\*\*** | **1** |
| ***Agathon ezoensis*** | **0.315** | **0.038 \*** | **1** | ***Epeorus nipponicus*** | **0.590** | **0.001 \*\*\*** | **1** |
|  |  |  |  | ***Glossosoma* spp.** | **0.588** | **0.001 \*\*\*** | **1** |
|  |  |  |  | *Antocha* spp. | 0.584 | 0.002 ** | 2 |
|  |  |  |  | *Ephemerella aurivillii* | 0.576 | 0.001 *** | 2 |
|  |  |  |  | **Orthocladiinae spp.** | **0.571** | **0.001 \*\*\*** | **1** |
|  |  |  |  | *Baetis* spp. | 0.531 | 0.002 ** | 2 |

<sup>a)</sup> The fish were classified into cold water species (< 20 °C) or cool-warm water species (20–30 °C) based on the thermal preference as reported by Natsumeda et al. (2010), Hatakeyama and Kitano (2018), and Suzuki et al. (2021). The macroinvertebrates were classified based on Arthington et al. (2006) and checked with the lists of Vieira et al. (2006) and Schmidt-Kloiber and Hering (2015). We classified macroinvertebrates into cold-water species (cold stenothermal or cool eurythermal: < 15 °C) and cool or warm water species (cool or warm eurythermal: 15–30 °C).

### 2.3. Role of watershed geology in creating climate-change refugia

#### 2.3.1. Target species

To test the contribution of watershed geology in creating climate-change refugia and their climate dependency, we predicted thermally suitable sites for the occurrence of lotic sculpins in the five study regions where water temperature modeling was undertaken. Lotic sculpins were found to be the most abundant fish in the mountain streams of the Sorachi and Chubu region from biotic surveys (see the Results section for details). Based on the difference in the distribution ranges of the target species, *Cottus nozawae* and *Cottus pollux* were used for the prediction in northern Japan (Okhotsk, Sorachi, Oshima) and central Japan (Kantou, Chubu), respectively. The occurrence of *C. nozawae* is predominantly restricted by the mean summer water temperature in northern Japan (Suzuki et al. 2021). Our previous study on the Sorachi River demonstrated that asymmetric gene flow of *C. nozawae* occurred from relatively low to high temperature streams, highlighting the potential importance of cold water tributaries in sustaining source populations (Nakajima et al. 2021). The indicator value analysis conducted for the present study showed that *C. nozawae* is the only indicator fish species for cold volcanic streams (see the Results section for details). *C. pollux* is endemic to eastern Japan, excluding the Hokkaido region, and prefers cold habitats (Natsumeda et al. 2010, Takeshita et al. 2017). However, no study has reported the thermal threshold for the occurrence like that of *C. nozawae*. We thus modeled the relationship of the occurrence probability for *C. pollux* to mean summer water temperature following Suzuki et al. (2021) by examining other candidate physiographic factors (see Fig. S4 details). Using the model to predict the mean summer water temperature, we predicted the future water temperature at each temperature monitoring site in northern and central Japan, respectively. We examined the difference in the change in the thermally suitable habitat for *C. nozawae* (< 16.1°C, Suzuki et al. 2021) in northern Japan and *C. pollux* (< 18.4 °C, Fig. S4) in central Japan between watershed geologies (non-volcanic vs. volcanic).

#### 2.3.2. Projection of future summer water temperatures

To predict future (2041–2060 and 2061–2080) changes in the thermally suitable habitat of *C. nozawae* in northern Japan (non-volcanic: 46 sites, volcanic: 31 sites) and *C. pollux* in central Japan (non-volcanic: 35 sites, volcanic: 28 sites), the future summer mean air temperature and the total precipitation (July–August) at each temperature monitoring site were obtained from the CHELSA global climate dataset. CHELSA is a high resolution (30 arc sec, approximately 1 km) global climate dataset based on mechanistic statistical downscaling of global ERA-Interim re-analysis data and future global circulation model outputs (Karger et al. 2017). We used the criterion suggested by Sanderson et al. (2015) to select a set of four global climate models (GCMs; MIROC5, MPI-ESM-MR, CESM1-CAM5, and IPSL-CM5A-MR) with a minimum interdependency to minimize the risk of spurious correlations among the GCMs, allowing for a better representation of the uncertainty in the climate model projections. We also selected the MRI-CGCM3 developed by the Japanese Meteorological Research Institute (Yukimoto et al. 2012) because it is a well-tested model which is commonly used for climate change predictions in Japan (Ishizaki et al. 2020). Future projected climates were calculated from the mean ensemble of the five GCMs for three contrasting Representative Concentration Pathways (RCPs): RCP2.6, RCP4.5, and RCP8.5. The contemporary summer water temperature at each site was calculated using contemporary climate data (1981–2010) with a 1 km grid resolution provided by the Japan Meteorological Agency.

## 3. Results

### 3.1. Influence of watershed geology on stream water temperature

Study streams from the middle to high latitudes in Japan had a high variety of mean summer water temperatures, which ranged from 7.7 to 22.2 °C. The climate at the monitoring site also had a high spatial variability ranging from 16.2 to 25.2 °C for the mean summer air temperature and 76 to 1,910 mm for the total summer precipitation (Fig. S5). Model averaging for three superior models showed five variables and two interaction terms as influencing factors for the summer water temperature (Table S2). The final model showed the variation in the mean summer water temperature with 0.62 of marginal R^2^, 0.96 of conditional R^2^, and 0.36 °C of RMSE (Table 2, Fig. 3). The mean summer water temperature was influenced by the mean summer air temperature, the total summer precipitation, the proportion of volcanic rocks, the site elevation and the proportion of agricultural land in the watershed. Cooling effects were related to the proportion of volcanic rocks in the watershed, the total summer precipitation, and site elevation. Warming effects were associated with the mean summer air temperature and the proportion of agricultural land in the watershed. The mean summer air temperature was the most influential, followed by the proportion of volcanic rocks in the watershed (Table 2, Fig.4). There was a significant interaction between the watershed geology and climate. The cooling effect due to an increase in volcanic rocks in the watershed was more pronounced in streams with less summer precipitation or lower summer air temperatures (Fig. 5). The interaction between the watershed geology and climates resulted in the regional difference in the geology-controlled cooling effect (i.e., the difference in the estimated mean summer water temperature between sites contributing catchments with volcanic and non-volcanic geological types in each region). The mean predicted cooling effects associated with the watershed geology in the Sorachi region, of which mean summer precipitation was 280 mm and mean summer air temperature was 18.6 °C, was 2.8 °C. The cooling effect in the Chubu region with 991 mm of mean summer precipitation and 21.4 °C of mean summer air temperature was 0.3 °C (Table S4).

**Table 2.**
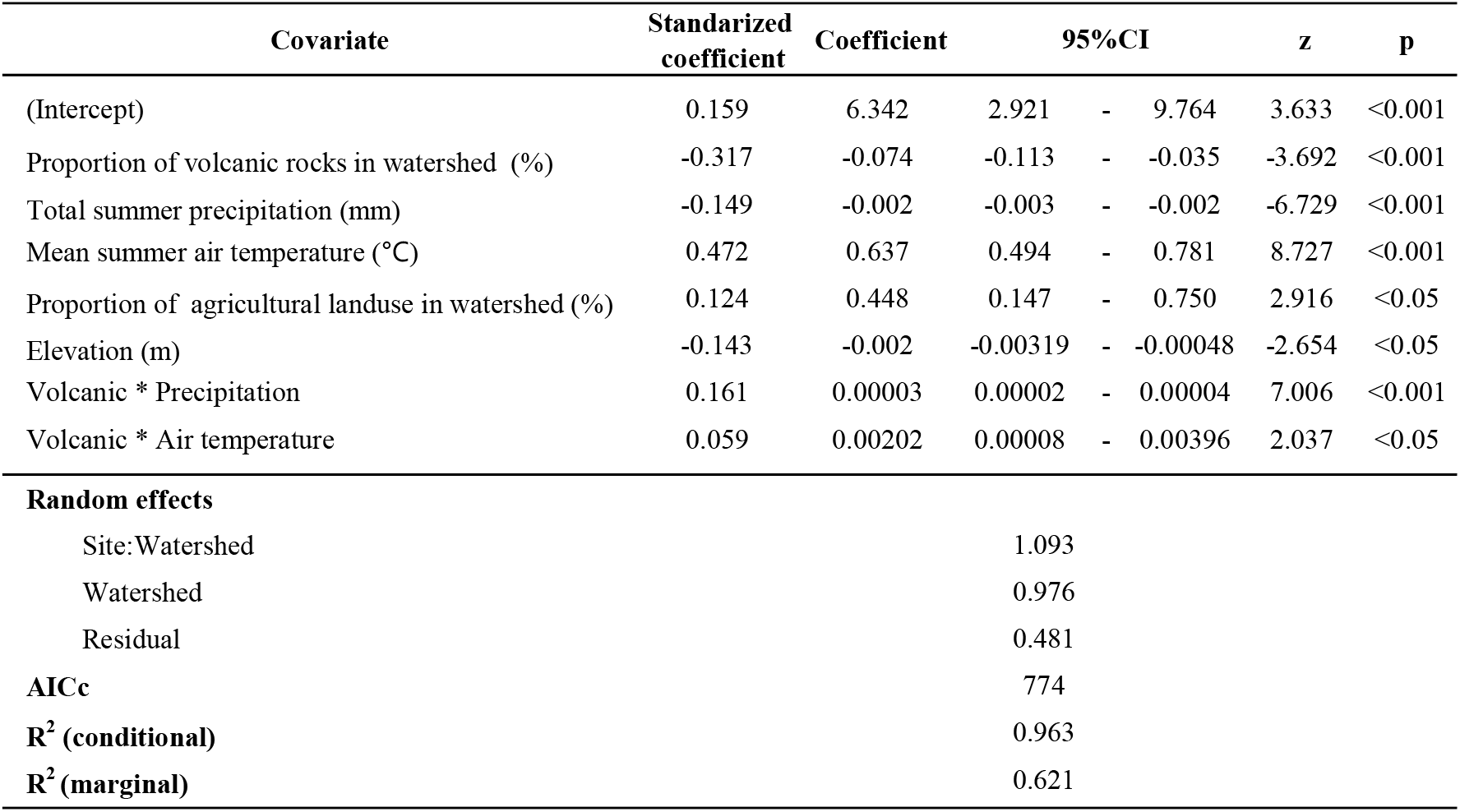
Summary of the results of the final model predicting the summer water temperature.

| Covariate | Standardized<br>coefficient | Coefficient | 95%CI |  | z | p |
| --- | --- | --- | --- | --- | --- | --- |
| (Intercept) | 0.159 | 6.342 | 2.921 | - 9.764 | 3.633 | <0.001 |
| Proportion of volcanic rocks in watershed (%) | -0.317 | -0.074 | -0.113 | - -0.035 | -3.692 | <0.001 |
| Total summer precipitation (mm) | -0.149 | -0.002 | -0.003 | - -0.002 | -6.729 | <0.001 |
| Mean summer air temperature (°C) | 0.472 | 0.637 | 0.494 | - 0.781 | 8.727 | <0.001 |
| Proportion of agricultural landuse in watershed (%) | 0.124 | 0.448 | 0.147 | - 0.750 | 2.916 | <0.05 |
| Elevation (m) | -0.143 | -0.002 | -0.00319 | - -0.00048 | -2.654 | <0.05 |
| Volcanic * Precipitation | 0.161 | 0.00003 | 0.00002 | - 0.00004 | 7.006 | <0.001 |
| Volcanic * Air temperature | 0.059 | 0.00202 | 0.00008 | - 0.00396 | 2.037 | <0.05 |
| **Random effects** |  |  |  |  |  |  |
| Site:Watershed |  |  | 1.093 |  |  |  |
| Watershed |  |  | 0.976 |  |  |  |
| Residual |  |  | 0.481 |  |  |  |
| **AICc** |  |  | 774 |  |  |  |
| **R<sup>2</sup> (conditional)** |  |  | 0.963 |  |  |  |
| **R<sup>2</sup> (marginal)** |  |  | 0.621 |  |  |  |

**Fig. 3.**
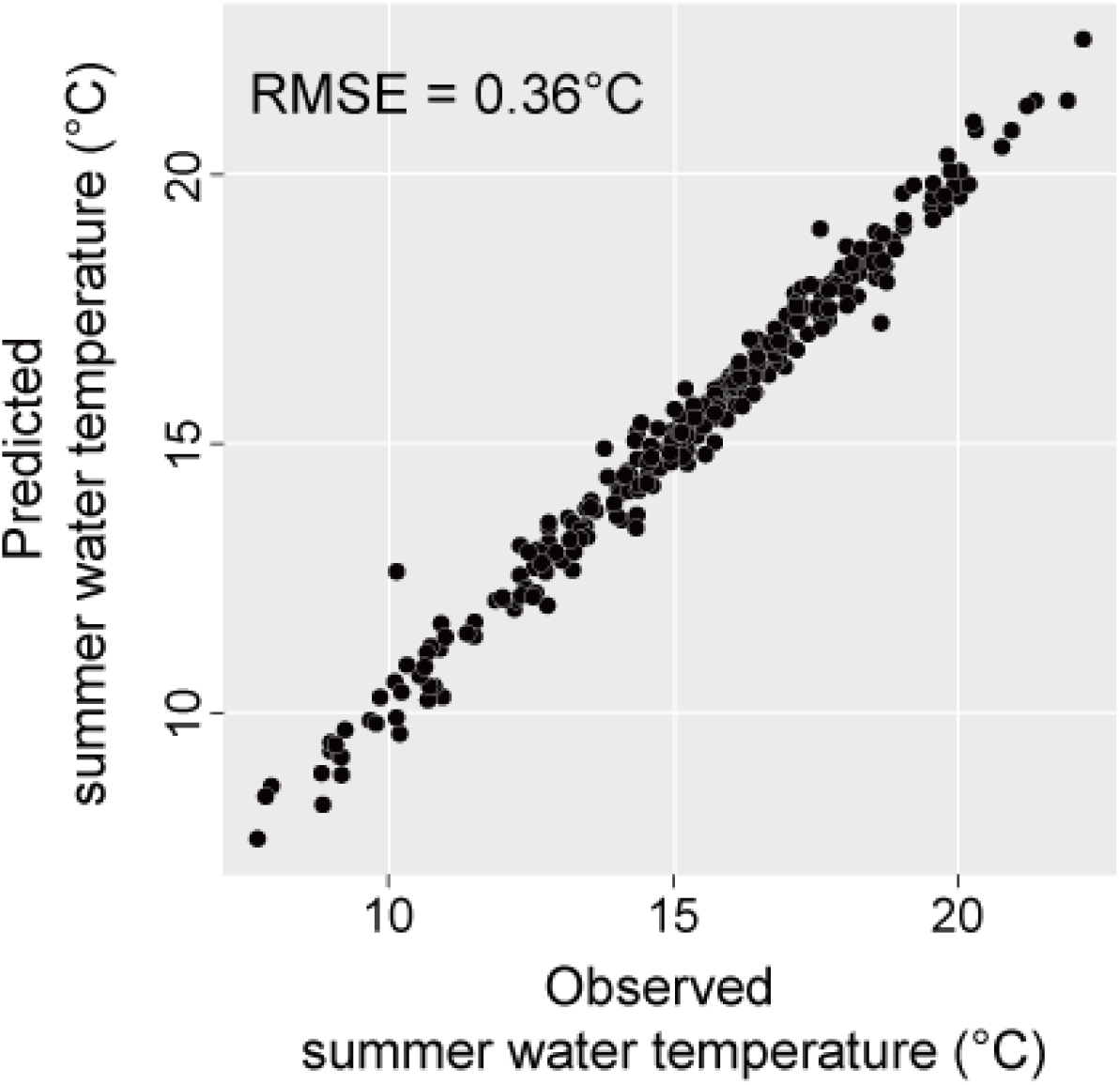
Scatter plot of the observed versus predicted mean summer water temperatures.

**Fig. 4.**
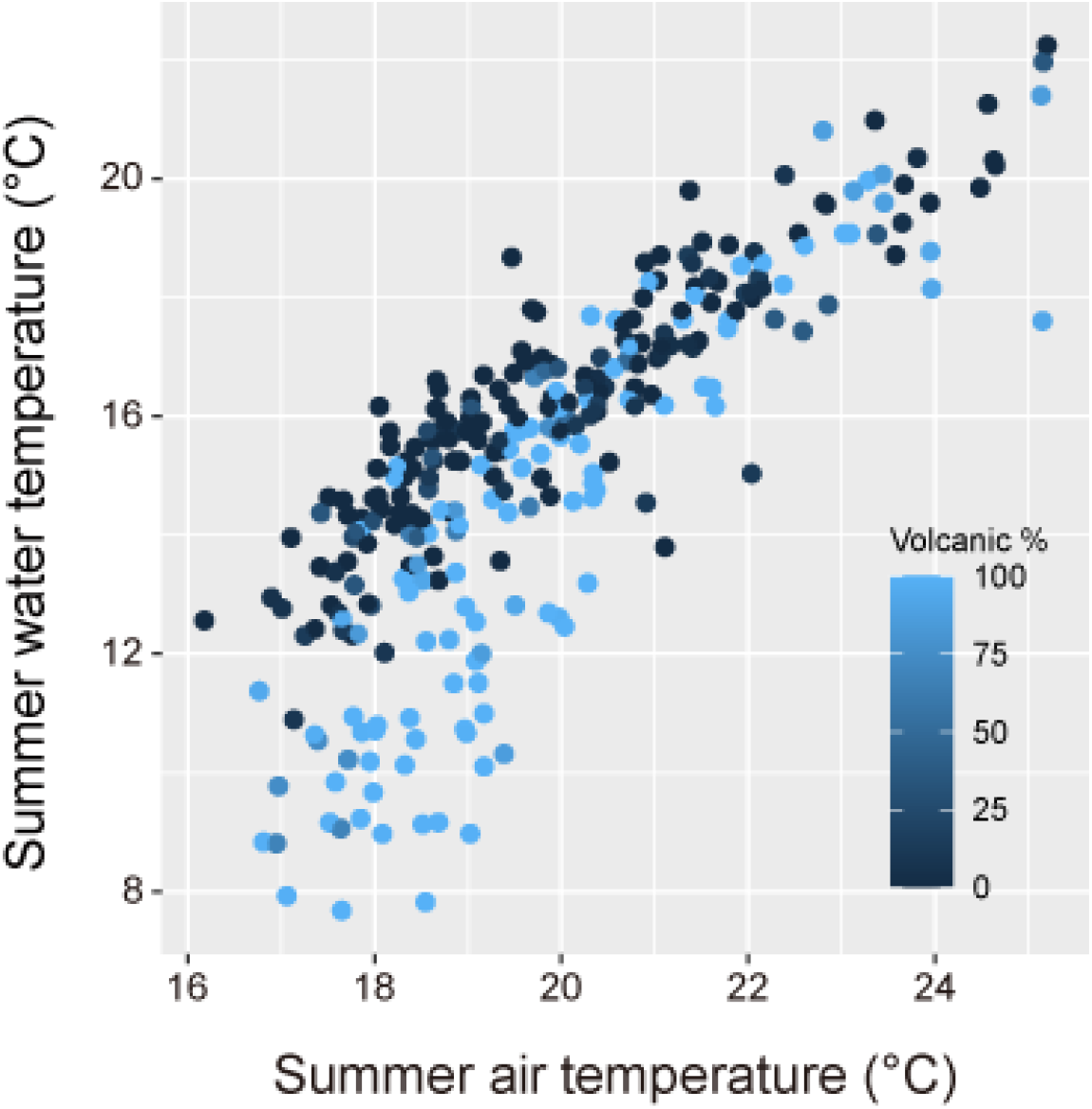
Relationship of the mean summer air temperature and watershed geology, namely the proportion of volcanic rock in the watershed, to the mean summer water temperature.

**Fig. 5.**
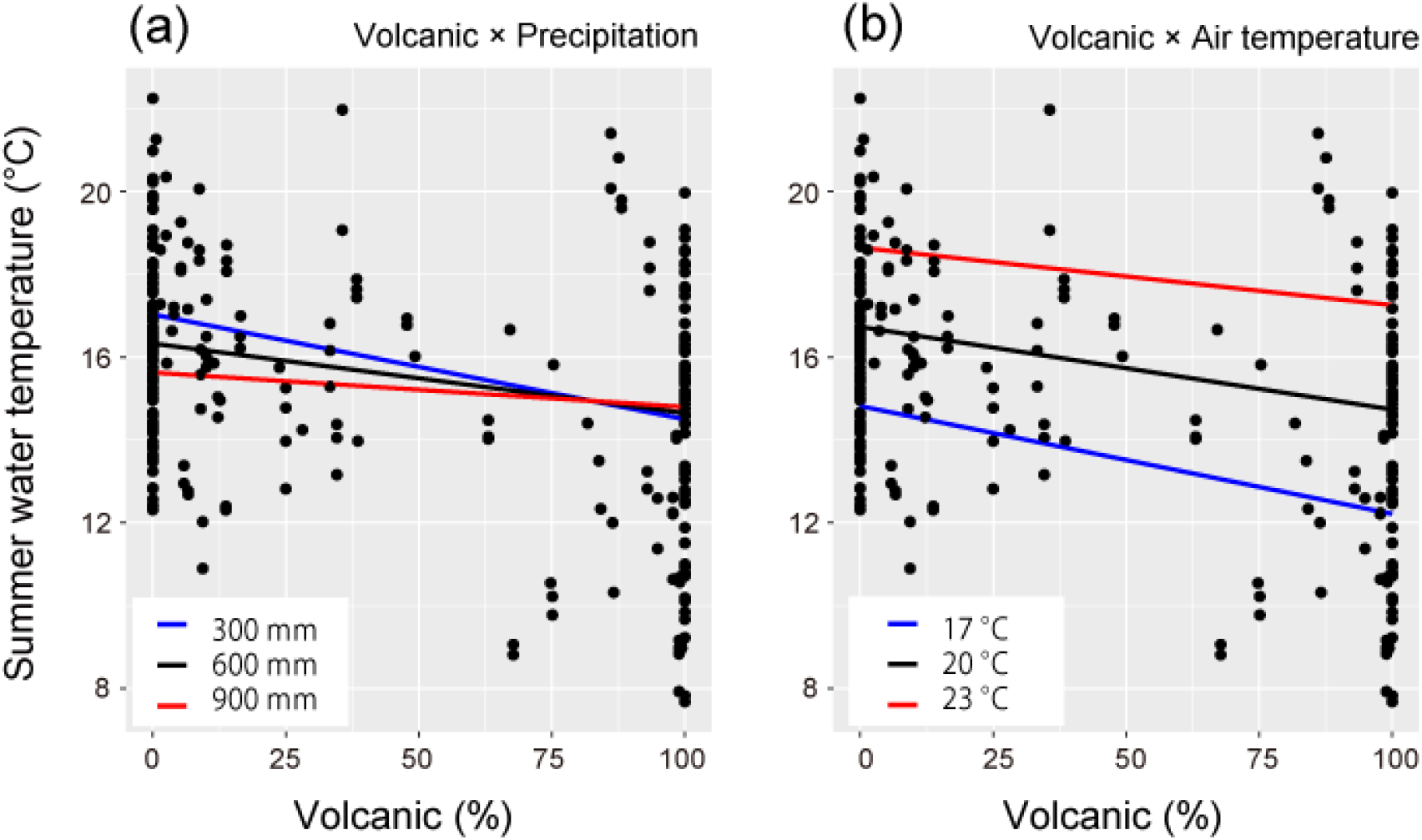
Relationship among the watershed geology, the mean summer water temperature, and climate. (a) The interactive effect between the watershed geology and the total summer recipitation, (b) the interactive effect between the watershed geology and the mean summer air temperature.

### 3.2. Response of stream community composition to watershed geology

In the Sorachi region, we observed 1,329 individuals from six fish species and 5,952 individuals from 60 macroinvertebrates (Table S3). The three most abundant fish species were *C. nozawae* (Scorpaeniformes, 1,040 inds.), *Barbatula* (Cypriniformes, 201 inds.), and *Salvelinus leucomaenis* (Salmoniformes, 73 inds.). The three most abundant macroinvertebrates were *Baetis* spp. (Ephemeroptera: 800 inds.), *Neophylax* spp. (Trichoptera, 201 inds.), and *Antocha* spp. (Diptera 73 inds.). In the Chubu region, we observed 2,110 individuals from 14 fish species and 8,600 individuals from 90 macroinvertebrates (Table S3). The three most abundant fish species were *C. pollux* (Scorpaeniformes, 821 inds.), *Rhinogobius* spp. (Cypriniformes, 336 inds.), and *Oncorhynchus masou ishikawai* (Salmoniformes, 278 inds.). The three most abundant macroinvertebrates were *Baetis themrmicus* (Ephemeroptera: 1,063 inds.), Orthocladiinae spp. (Diptera, 629 inds.), and *Simulium spp*. (Diptera 456 inds.).

In the Sorachi region where there was low summer precipitation and low air temperatures, fish and macroinvertebrate community composition differed significantly between the geological types (Fig. 6a, b). The stress value for the ordination of each taxon, which is an index of the fit of the reproduced similarity matrix to the observed similarity was 0.10 (fish) and 0.18 (macroinvertebrates), suggesting a good representation (Clarke 1993). For both taxa, the proportion of volcanic rocks in the watershed (fish: *r*^2^ = 0.43, *p* = 0.002; macroinvertebrates: r^2^ = 0.51, *p* = 0.0002) and the mean summer water temperature (fish: *r*^2^ = 0.56, *p* = 0.0004; macroinvertebrates: r^2^ = 0.63, *p* = 0.0002) were significantly correlated with community composition (Fig. 6a, b). Indicator value analyses showed that stream communities in volcanic streams were characterized by cold-water species and vice versa (Table 3). *B. barbatula* (cool-water species) is a representative species of non-volcanic stream communities, and *C. nozawae* (cold-water species) is the sole species representing volcanic stream communities. In the macroinvertebrate community, five and 18 species were confined to non-volcanic and volcanic streams, respectively. For the non-volcanic streams, two cold-water species were included among the five representative species, but the fidelity values were not high (≤ 0.4). In the volcanic streams, species with high fidelity values were dominated by cold-water species. In the Chubu region with high levels of summer precipitation and high air temperatures, the stress value for the ordination of fish and macroinvertebrates showed a good fit with 0.12 and 0.16, respectively (Fig. 6c, d). As well as the community-environment relationship found in the Sorachi region, the fish and the macroinvertebrate community composition were found to be significantly associated with the mean summer water temperature (fish: *r*^2^ = 0.65, *p* = 0.0002; macroinvertebrates: r^2^ = 0.78, *p* = 0.0002). However, the stream community composition in the Chubu region showed no significant difference between geological types in contrast with those of the Sorachi region (Fig. 6c, b).

**Fig. 6.**
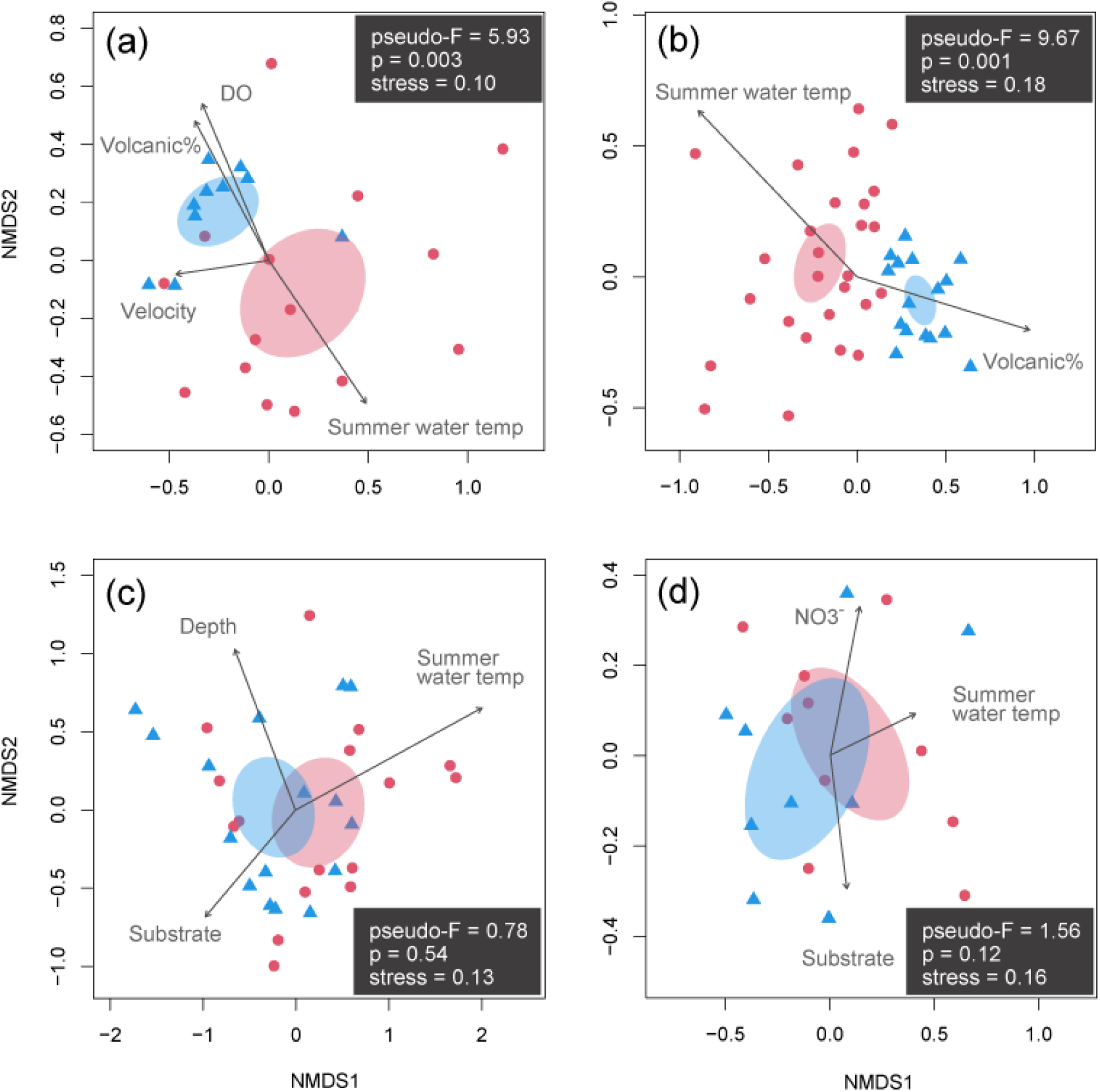
NMDS ordination plots for each geological type and significantly correlated local environmental factors. (a) Fish species composition in the Sorachi, (b) macroinvertebrate species composition in the Sorachi, (c) fish species composition in the Chubu and (d) macroinvertebrate species composition in the Chubu. Blue circles and red triangles indicate volcanic and non-volcanic streams, respectively. The length of the arrows shows the correlation between each environment and the community composition. Ellipses were drawn at the 95% level of confidence for each geological type. Results of PERMANOVA, pseudo-F and P-value, are also shown in each panel.

### 3.3. Thermally suitable habitat for lotic sculpins under climate change

Under contemporary climatic conditions (1981–2010), both geological types moderately sustain thermally suitable sites for lotic sculpins. In northern Japan, the ratios for *C. nozawae* (< 16.1 °C) in non-volcanic and volcanic streams were 78.2 and 96.8%, respectively. For *C. pollux* (< 18.4 °C) in central Japan, the ratios for non-volcanic and volcanic streams were 37.1 and 71.4%, respectively.

By 2041–2060, the predicted mean increase in the summer water temperature across sites under RCPs 2.6, 4.5, and 8.5 were 1.4, 1.7, and 2.1 °C in northern Japan, and 2.1, 2.4, and 2.7 °C in central Japan, respectively. The mean summer water temperature was predicted to increase further by 2061–2080 in both geological types in both regions (Fig. 7). As a result of this warming trend, thermally suitable habitats for sculpins are projected to decrease. However, the predicted temporal changes greatly differ between geological types and regions (Fig. 7). In northern Japan with a high geology-controlled cooling effect, the temporal pattern of thermal habitat loss for *C. nozawae* differed substantially between geological types (Fig. 7a). The thermally suitable habitats in non-volcanic streams will decline under all emission scenarios, with 8.7% of sites remaining thermally suitable for *C. nozawae* under RCP 8.5 in 2061–2080. In contrast, volcanic streams in northern Japan will remain cooler than non-volcanic streams, resulting in a much higher proportion of sites remaining thermally suitable for the species by 2061–2080 (83.9% and 74.2% respectively for RCPs 2.6 and 4.5). However, by 2061–2080, conditions under RCP 8.5 are projected to moderately deteriorate for the species even in sites located in volcanic streams (61.3%). Contrary to the projections for northern Japan, thermally suitable habitat for *C. pollux* in central Japan with a low geology-controlled cooling effect will largely disappear regardless of the geological type. Streams in both geological types will lose thermally suitable habitats under all emission scenarios, with less than 25% of sites remaining thermally suitable under even the lowest emission scenario, RCP 2.6. These predictions based on the mean ensemble model projections generally hold for projections based on individual climate models (Fig. S6,7). The scenario analysis results highlight the importance of geology-controlled climate-change refugia in mountain streams and their climate dependency.

**Fig. 7.**
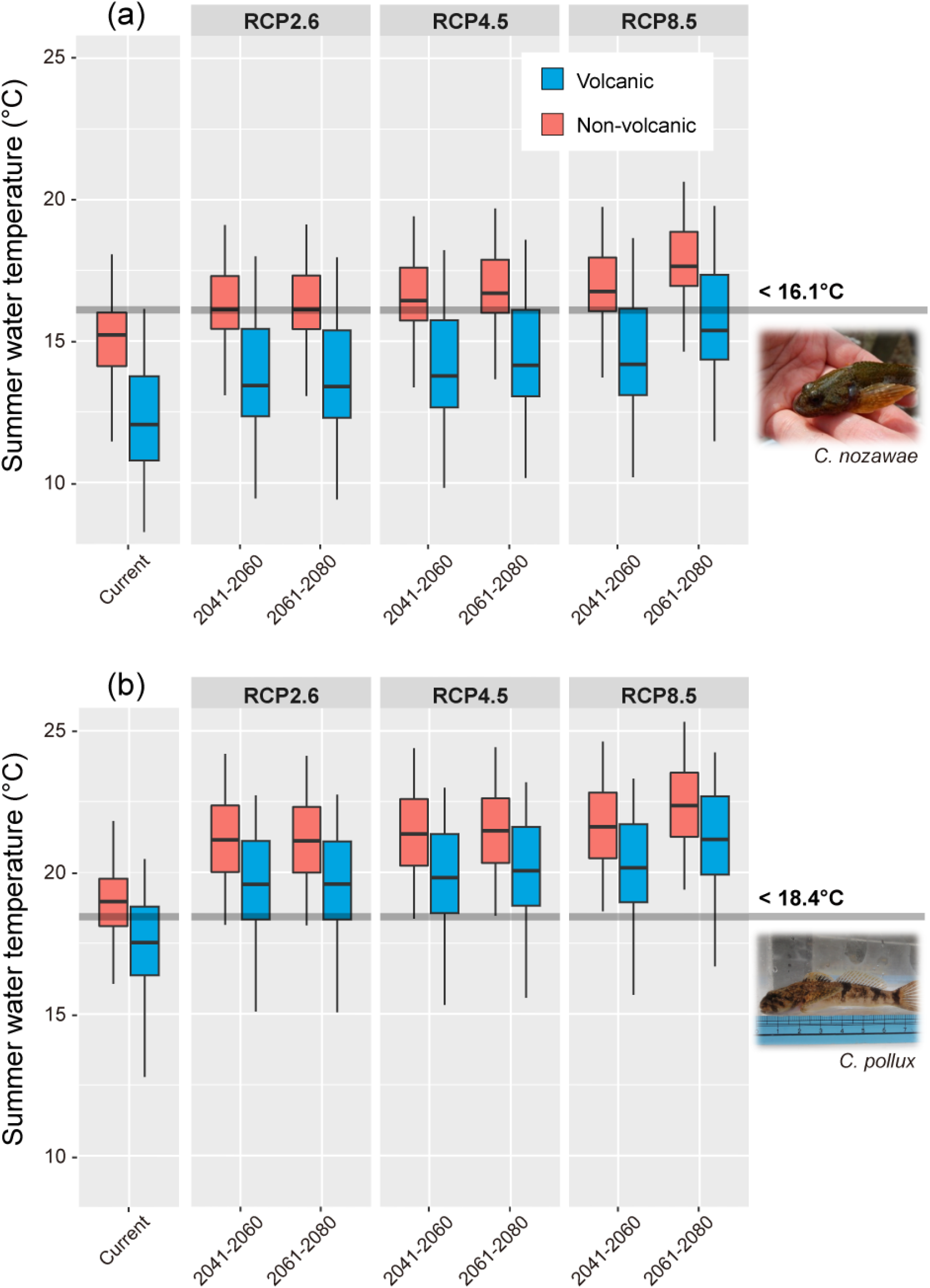
The current and future mean summer water temperature and the relation to the thermal threshold for lotic sculpins in a) northern Japan and b) central Japan. Predictions are shown by each period and the RCP. The lines in the middle of the boxes represent the median, and the lower and upper ends of the boxes are the 25 and 75% quartiles, respectively.

## 4. Discussion

In the present study, we modeled the mean summer water temperature of Japanese mountain streams by considering the interaction between the watershed geology and climatic conditions in addition to their independent effects. The cooling effect associated with watershed geology was predicted to create climate change refugia for cold water species. The distribution shift to upstream is a key ecological response to increased thermal stress in stream ecosystems (e.g., Comte and Grenouillet 2013, Isaak and Rieman 2013, Herrera - R et al. 2020). In this context, high elevation mountain streams with cooler habitats are considered climate change refugia (Isaak et al. 2016). The conventional refugia are also supported by the present study because summer water temperature varies with altitudinal gradient (Fig. S8). Our findings indicate that spatial variation in the underlying geology is vital in informing the distribution of climate change refugia as well as the altitudinal gradient. Given the impacts of global warming and habitat fragmentation on freshwater ecosystems (Scheffers et al. 2016, Ishiyama et al. 2018, Herrera-R et al. 2020), connectivity restoration and conservation between current and future suitable habitats is one of the key adaptation measures. Further studies are needed to deepen our understanding of the underlying mechanisms of the interplay between watershed geology and climate, but findings from the present study highlight that we should consider watershed geology, climate variability, and the interaction among those covariates for effective connectivity management in mountain streams exposed to future climate change.

### 4.1. Stream summer water temperature along a geological and climatic gradient

Our statistical approach revealed that the proportion of volcanic rocks in the watershed determines the stream summer water temperature following air temperature. Tague et al. (2007) showed that the stream summer water temperature in spring-fed streams was colder than that in shallow subsurface flow-fed streams. The flow path distinction relates to the bedrock permeability; water infiltrating into the soil mainly discharges into headwater streams as shallow subsurface flow in catchments with less permeable bedrock whereas it leaves the headwaters as groundwater flow within bedrock and discharges downstream as springs in catchments with highly permeable bedrock (Tague et al. 2007, Richardson et al. 2020). Comparison studies of watersheds in Japan at the 10^1^–10^2^ km^2^ scale showed that Quaternary and Tertiary volcanic watersheds had greater low-flow specific discharges, suggesting a higher bedrock permeability than sedimentary watersheds (Shimizu 1980, Mushiake et al. 1981). Geographic source separation of the baseflow stream water at multiple points also revealed that specific discharge of not only deep groundwater but also shallow groundwater increased with the drainage area in Japanese volcanic watersheds dominated by andesite (i.e., non-alkaline mafic volcanic rocks), while that of shallow groundwater was almost constant regardless of the area in sedimentary rock watersheds (Iwasaki et al. 2021). Based on this result and that of previous studies, they presented the presumed mechanisms by which shallow groundwater can discharge downstream beyond the drainage divide because of the flow path within thick porous andesite lava at volcanic watersheds whereas it discharges to the headwater streams because of the limited flow path within the soil and the thin weathered bedrock layer at the sedimentary watersheds. The flow mechanism fits with our study systems because the dominant volcanic lithology in Iwasaki et al. (2021) is the same as that of our study. Following the non-alkaline mafic volcanic geology, the non-alkaline pyroclastic flow volcanic rocks, such as clastic pumice, are the second most common volcanic lithology in the watersheds in this study. There are many springs and groundwater-fed streams with past pyroclastic flows in the Japanese region (Inoue et al. 2009, Kawai et al. 2013). This is likely because pyroclastic flow layers have high permeability with groundwater movement through a bedrock layer, including thick deposits of volcanic materials (Fujimoto et al. 2016). Due to the high permeability, the rich groundwater inputs downstream would have contributed to the decrease in the summer water temperature in volcanic streams.

More importantly, the cooling effect of volcanic rocks depends on the climatic conditions. A strong cooling effect was observed in the watersheds with low summer precipitation (Fig. 5a). During rainfall, overland or subsurface flow pathways create hydrograph peaks, even in catchments underlain by permeable volcanic bedrock (Iwagami et al. 2010, Muñoz-Villers and McDonnell 2012). This flow-regime pattern indicates that the contribution of cold groundwater inputs to stream discharge decreases with increasing precipitation during the summer. Therefore, the water temperature response to watershed geology varies with the total summer precipitation (i.e., the relative contribution of groundwater input). Thereafter, the influence of watershed geology on the summer water temperature was diluted by an increased summer air temperature (Fig. 5b). This is likely due to the increased temperature of groundwater sources. Groundwater pathways are categorized into shallow or deep types, depending on the depth of the aquifer source (Hare et al. 2021). Unlike deep groundwater, groundwater flowing near the surface is easily affected by changes in the land surface temperature (Kurylyk et al. 2013). In this study, in watersheds with a high air temperature, the cooling effect associated with shallow groundwater may be weakened by increased land-groundwater heat transport, resulting in the interaction between watershed geology and the summer air temperature.

### 4.2. Geology controlled climate-change refugia

As we expected, stream community composition was found to reflect the underlying geology. Fish and macroinvertebrates community composition in the Sorachi region differed between the non-volcanic and volcanic streams. This is predominantly controlled by the difference in the summer water temperature (Fig. 6a, b). Further indicator species analyses showed that the volcanic stream hosted communities associated with more cold-water species. Rising water temperature can lead to negative ecological impacts, including reduced food consumption (Selong et al. 2001), swimming ability or equilibrium (Dallas and Rivers-Moore 2012, Yamada et al. 2020), decreased fecundity (Brittain 1991), and ultimately mortality. These thermal stresses are more evident in cold-water species. We selected natural mountain streams with low levels of human disturbance (see “Biotic survey” for details). Therefore, the community assembly pattern related to the watershed geology may reflect natural thermal filtering. As well as results from the Sorachi region, the mean summer water temperature was the influential factor for the fish and macroinvertebrate community composition in the Chubu region. However, the stream community composition in the Chubu region showed no significant difference between geological types. The difference in the importance of watershed geology in shaping stream communities may be attributed to the climate-dependent cooling effect of groundwater inputs. The stronger cooling effect of volcanic rocks was observed in watersheds with less summer precipitation and higher air temperatures. The amount of summer precipitation and the air temperature in the Chubu region was considerably higher compared to the Sorachi region (Table 1) that exhibited a significant influence of watershed geology on the stream community composition. The other candidate factors selected in the water temperature modeling, such as climate, land use and elevation, would indirectly shape stream communities along the thermal gradients in the Chubu region.

Given the continuous impacts of global warming, streams providing cold habitats will be more important as climate-change refugia. Scenario analyses for northern Japan have predicted that many thermally suitable habitats for *C. nozawae* will remain in volcanic streams compared with non-volcanic streams especially for the low (RCP 2.6) and middle (RCP 4.5) emission scenarios. This prediction highlights the importance of volcanic watersheds with high groundwater contributions in creating climate-change refugia for cold-water species. However, geology-controlled refugia do not occur everywhere and depend on climatic conditions as shown in the constructed water temperature models and scenario analyses for central Japan. In regions where future climates show projected high precipitation and air temperatures, the geological contribution to climate-change refugia would be reduced. In streams where movement is constrained within the network, headwaters at higher elevation provide one of the most viable and accessible options for aquatic species tracking their shifting thermal niches as many previous studies have demonstrated (Comte and Grenouillet 2013, Isaak and Rieman 2013). However, geology-controlled thermal heterogeneity and its influence on stream communities are likely more related to the concept of climate-refugia as patches of suitable habitat embedded within larger areas or landscapes with unsuitable conditions. Previous studies and our findings suggest that both types of cold refugia are complementary and should be considered together when designing watershed-scale management strategies for biodiversity conservation in response to climate change.

The identification of climate-change refugia has gained growing attention in the context of climate change adaptation (Morelli et al. 2016, Morelli et al. 2020). However, direct measurement of groundwater inputs at broad spatial scales for mapping climate-change refugia is difficult, even though although climate-change adaptation measures are needed across a wide range of regions and continents. Some indices measuring the contribution of groundwater inputs in the statistical prediction of stream summer water temperature have already been used. Previous models account for the influence of groundwater on thermal sensitivity from the interactions of the air-water temperature being measured (Snyder et al. 2015) and the base flow index (BFI) based on measured hydrographs (Isaak et al. 2017). Substitutes for groundwater input improved predictions, which have a practical limitation because thermal or flow regime data are required for predicting the water temperature at target streams. In this study, we used the proportion of the volcanic rocks in the watershed as the surrogate and constructed a summer stream temperature model with accuracy (RMSE = 0.36 °C, Fig. 3). The model performance improved with the interplay of the climate conditions. Both climate (e.g., CHELSA) and geological maps (e.g., OneGeology) are readily available at broad spatial scales. Therefore, coarse filtering based on the watershed geology and climate can be a straightforward approach for inferring the influence of groundwater input which varies with climate variability, contributing to the identification of climate-change refugia in mountain streams at various spatiotemporal scales. However, current associations between the geological surrogate and the water temperature can no longer hold for the future under accelerated climate change, as these relationships are not time invariant and depend on how climate change including air temperature, rainfall, and snowmelt alter the recharge, routing and discharge of groundwater into streams. Further examination using a process-based approach that considers both atmospheric and land surface processes, such as a distributed hydrothermal model, are needed to confirm the importance of watershed geology as abiotic surrogate and its temporal universality.

## 5. Conclusions

Following the identification of refugia, an important management action is to reconnect climate-change refugia to each other and nearby non-refugia habitats to improve long-term access to refugial areas (Morelli et al. 2016). Our findings highlight the importance of considering the spatial variation in the watershed geology in connectivity management. In the case of *C. nozawae* in norther Japan (Fig. 7a), populations in non-volcanic streams will be exposed to severe thermal stress in the near future, 2041–2060, even under the lowest emission scenario. The species-distribution shift along the elevational gradient can also fail in non-volcanic streams. For example, most of the monitoring sites in the Sorachi river, ranging from 78 to 520 m in altitude, will exceed the thermal threshold for *C. nozawae* (Fig. S8). Considering these trends, the management priority should be set into hydrologic connectivity between non-volcanic streams and nearby volcanic streams as refugia to ensure the persistence of these threatened populations. This type of connectivity management would also effectively conserve cold-water species that have high mobility. This is because migratory cold-water species such as salmonids can seasonally use the thermal heterogeneity in dendritic networks and habitats that are warm during the summer and provide the majority of growth potential during other seasons (Armstrong et al. 2021). All volcanic streams do not function equally as thermal refugia in changing climates. In the predictions for northern Japan, volcanic streams will serve as climate-change refugia until the middle emission scenario. However, approximately half of them will exceed the thermal threshold for the occurrence of *C. nozawae* by 2061–2080 under the most severe scenario RCP 8.5 (Fig. 7a). Therefore, connections among volcanic streams (i.e., elevational shifts in species distribution) should also be considered in the next phase of climate-change refugia management to ensure a distribution shift to more suitable volcanic streams and sufficient spatial extent in the available thermal habitat under future climate conditions. In this way, climate-change adaptation considering watershed geology can inform more effective conservation of cold-water species. However, managers should also remember that the contribution of watershed geology in creating climate-change refugia would not be universal because thermal heterogeneity in the mountain streams was affected by the interaction between geology and climate as well as these independent effects. Contrary to projections for northern Japan, thermally suitable habitat for *C. pollux* in central Japan was projected to be largely lost regardless of the geological type. Our stream temperature predictive models and scenario analyses indicate that the geology-controlled climate-change refugia should be paid more attention in regions with lower precipitation or air temperature.

In this study, we classified the Earth’s subsurface rock lithology into two types (non-volcanic vs. volcanic) because of the limited temperature monitoring network. Although our models showed good predictions for the summer water temperatures of mountain streams, the geological classification we used had limitations in generalizing the usefulness of watershed geology as a surrogate for groundwater contribution. Some previous studies reported that younger volcanic bedrock has higher permeability than older volcanic bedrock (Mushiake et al. 1981, Tague and Grant 2004). Similar differences in the flow regime were also found between specific non-volcanic lithologies, namely between accretionary complex and plutonic rocks (Onda et al. 2006). These studies suggest a potential difference in the amounts or pathways of groundwater inputs depending on the geological age and type, which were not considered in the present study. Further studies covering a variety of geological ages and types based on a more extensive monitoring network and integration with species responses with a variety of traits will promote a better understanding of the role of nature’s stage in global warming adaptation.

## Acknowledgments

We thank Suzuki Kaiji and Todate Shun for their help in conducting the field and laboratory work. This research was funded by JSPS KAKENHI 18K18221, JSPS KAKENHI 19H04314, the research fund for the Ishikari and Tokachi Rivers provided by the Ministry of Land, Infrastructure, Transport, and Tourism (MLIT) of Japan and the Environment Research and Technology Development Fund grant (JPMEERF20202004) of the Environmental Restoration and Conservation Agency of Japan.

