## Supplementary materials for "The role of geology in creating stream climate-change refugia along climate gradients"

**Ishiyama Nobuo, Sueyoshi Masanao, García Molinos Jorge, Iwasaki Kenta, Negishi N Junjiro, Koizumi Itsuro, Nagayama S, Nagasaka Akiko, Nagasaka Yu, Nakamura Futoshi**

**Corresponding author**

Nobuo Ishiyama:

**Table S1** Summary of environments for NDMS analyses.

| Environments for NMDS analyses | Mean ± SD | Environments for NMDS analyses | Mean ± SD |
|---|---|---|---|
| **Sorachi** | | **Chubu** | |
| **(Fish)** | | **(Fish)** | |
| Mean summer water temperature (°C) | 12.7 ± 2.5 | Mean summer air temperature (°C) | 17.8 ± 2.2 |
| Volcanic rocks in the watershed (%) | 45.7 ± 43.4 | Volcanic rocks in the watershed (%) | 54.8 ± 42.2 |
| Mean water depth (m) | 0.21 ± 0.05 | Mean water depth (m) | 0.36 ± 0.11 |
| Mean current velocity (m/s) | 0.75 ± 0.19 | Mean current velocity (m/s) | 0.38 ± 0.16 |
| Substrate coarseness | 3.6 ± 0.2 | Substrate coarseness | 3.5 ± 0.6 |
| Dissolved oxygen (mg/l) | 10.7 ± 0.6 | Dissolved oxygen (mg/l) | 9.7 ± 1.1 |
| **(Macroinvertebrates)** | | **(Macroinvertebrates)** | |
| Mean summer water temperature (°C) | 13.5 ± 2.6 | Mean summer air temperature (°C) | 16.7 ± 3.5 |
| Volcanic rocks in the watershed (%) | 42.2 ± 44.6 | Volcanic rocks in the watershed (%) | 49.9 ± 44.7 |
| Mean water depth (m) | 0.18 ± 0.04 | Mean water depth (m) | 0.22 ± 0.06 |
| Mean current velocity (m/s) | 0.66 ± 0.16 | Mean current velocity (m/s) | 0.41 ± 0.11 |
| Substrate coarseness | 3.1 ± 0.3 | Substrate coarseness | 3.8 ± 0.4 |
| $NO^{3-}$ (mg/l) | 1.9 ± 3.3 | $NO3^{-}$ (mg/l) | 0.82 ± 0.80 |

**Table S2** Results of the model selection and averaging.

| Rank | Air | Area | Elev | Agri | Precipi | Slope | Vol | Vol*Air | Vol*Preicipi | df | AICc | delta AICc | Akaike weight |
|---|---|---|---|---|---|---|---|---|---|---|---|---|---|
| 1 | 0.64 | | −0.0018 | 0.45 | −0.0023 | | −0.07 | 0.0020 | 0.000028 | 11 | 774.30 | 0.00 | 0.35 |
| 2 | 0.70 | | −0.0019 | 0.46 | −0.0023 | | −0.03 | | 0.000027 | 10 | 776.23 | 1.92 | 0.14 |
| 3 | 0.64 | | −0.0018 | 0.46 | −0.0023 | 0.01 | −0.07 | 0.0020 | 0.000028 | 12 | 776.31 | 2.00 | 0.13 |
| 4 | 0.64 | 0.00 | −0.0018 | 0.45 | −0.0023 | | −0.07 | 0.0020 | 0.000028 | 12 | 776.48 | 2.18 | 0.12 |
| 5 | 0.70 | | −0.0019 | 0.47 | −0.0023 | 0.01 | −0.03 | | 0.000027 | 11 | 778.23 | 3.92 | 0.05 |
| **95% CIs** | | | | | | | | | | | | | |
| 2.5% | 0.50 | | −0.00322 | 0.15 | −0.00301 | −0.04 | −0.11 | 0.00007 | 0.000020 | | | | |
| 97.5% | 0.80 | | −0.00049 | 0.76 | −0.00164 | −0.06 | −0.02 | 0.00397 | 0.000036 | | | | |

**Table S3** List of analyzed species in the NMDS.

| Order | Family | Genus | Species |
|---|---|---|---|
| | | **Sorachi** | |
| (Fish) | | | |
| Salmoniformes | Salmonidae | Salvelinus | *Salvelinus leucomaenis leucomaenis* |
| Salmoniformes | Salmonidae | Salvelinus | *Salvelinus malma krascheninnikovi* |
| Salmoniformes | Salmonidae | Oncorhynchus | *Oncorhynchus mykiss* |
| Perciformes | Cottidae | Cottus | *Cottus nozawae Snyder* |
| Cypriniformes | Cobitidae | Barbatula | *Barbatula barbatula* |
| Petromyzontiformes | Petromyzontidae | Lethenteron | *Lethenteron* sp. *N* |
| (Macroinvertebrates) | | | |
| Ephemeroptera | Leptophlebiidae | Paraleptophlebia | *Paraleptophlebia japonica* |
| Ephemeroptera | Ephemeridae | Ephemera | *Ephemera* spp. |
| Ephemeroptera | Ephemerellidae | Cincticostella | *Cincticostella elongatula* |
| Ephemeroptera | Ephemerellidae | Cincticostella | *Cincticostella nigra* |
| Ephemeroptera | Ephemerellidae | Cincticostella | *Cincticostella orientalis* |
| Ephemeroptera | Ephemerellidae | Drunella | *Drunella basalis* |
| Ephemeroptera | Ephemerellidae | Drunella | *Drunella ishiyamana* |

| | | | |
|---|---|---|---|
| Ephemeroptera | Ephemerellidae | Drunella | *Drunella sachalinensis* |
| Ephemeroptera | Ephemerellidae | Drunella | *Drunella trispina* |
| Ephemeroptera | Ephemerellidae | Ephemerella | *Ephemerella aurivillii* |
| Ephemeroptera | Ephemerellidae | Teleganopsis | *Teleganopsis punctisetae* |
| Ephemeroptera | Ameletidae | Ameletus | *Ameletus* spp. |
| Ephemeroptera | Baetidae | Baetis | *Baetis* spp. |
| Ephemeroptera | Heptageniidae | Cinygmula | *Cinygmula* spp. |
| Ephemeroptera | Heptageniidae | Epeorus | *Epeorus latifolium* |
| Ephemeroptera | Heptageniidae | Epeorus | *Epeorus nipponicus* |
| Ephemeroptera | Heptageniidae | Rhithrogena | *Rhithrogena japonica* |
| Plecoptera | Nemouridae | Amphinemura | *Amphinemura* spp. |
| Plecoptera | Nemouridae | Nemoura | *Nemoura* spp. |
| Plecoptera | Nemouridae | Protonemura | *Protonemura* spp. |
| Plecoptera | Chloroperlidae | | *Chloroperlidae* spp. |
| Plecoptera | Perlidae | Gibosia | *Gibosia* spp. |
| Plecoptera | Perlidae | Kamimuria | *Kamimuria quadrata* |
| Plecoptera | Perlidae | Kamimuria | *Kamimuria tibialis* |
| Plecoptera | Perlidae | Kamimuria | *Kamimuria* spp. |
| Plecoptera | Perlodidae | | Perlodidae spp. |

| | | | |
|---|---|---|---|
| Trichoptera. | Hydropsychidae | Cheumatopsyche | *Cheumatopsyche infascia* |
| Trichoptera. | Hydropsychidae | Hydropsyche | *Hydropsyche orientalis* |
| Trichoptera. | Philopotamidae | Dolophilodes | *Dolophilodes* spp. |
| Trichoptera. | Polycentropodidae | Plectrocnemia | *Plectrocnemia* spp. |
| Trichoptera. | Stenopsychidae | Stenopsyche | *Stenopsyche marmorata* |
| Trichoptera. | Glossosomatidae | Glossosoma | *Glossosoma* spp. |
| Trichoptera. | Hydrobiosidae | Apsilochorema | *Apsilochorema sutshanum* |
| Trichoptera. | Rhyacophilidae | Rhyacophila | *Rhyacophila* spp. |
| Trichoptera. | Apataniidae | Apatania | *Apatania* spp. |
| Trichoptera. | Lepidostomatidae | Brachycentrus | *Brachycentrus americanus* |
| Trichoptera. | Lepidostomatidae | Micrasema | *Micrasema hanasensis* |
| Trichoptera. | Lepidostomatidae | Lepidostoma | *Lepidostoma* spp. |
| Trichoptera. | Limnephilidae | Dicosmoecus | *Dicosmoecus jozankeanus* |
| Trichoptera. | Limnephilidae | Hydatophylax | *Hydatophylax festivus* |
| Trichoptera. | Uenoidae | Neophylax | *Neophylax* spp. |
| Diptera | Pediciidae | Dicranota | *Dicranota* spp. |
| Diptera | Limoniidae | Antocha | *Antocha* spp. |
| Diptera | Limoniidae | Hexatoma | *Hexatoma* spp. |
| Diptera | Limoniidae | | Limoniidae spp. |

| | | | |
|---|---|---|---|
| Diptera | Tipulidae | Tipulidae | Tipulidae spp. |
| Diptera | Blephariceridae | Agathon | *Agathon japonicus* |
| Diptera | Blephariceridae | Agathon | *Agathon ezoensis* |
| Diptera | Blephariceridae | Bibiocephala | *Bibiocephala infuscata infuscata* |
| Diptera | Psychodidae | Pericoma | *Pericoma* spp. |
| Diptera | Ceratopogonidae | | Ceratopogonidae spp. |
| Diptera | Chironomidae | Tanypodinae | Tanypodinae spp. |
| Diptera | Chironomidae | Diamesinae | Diamesinae spp. |
| Diptera | Chironomidae | Orthocladiinae | Orthocladiinae spp. |
| Diptera | Chironomidae | Chironominae | Chironominae spp. |
| Diptera | Simuliidae | Prosimulium | *Prosimulium* spp. |
| Diptera | Simuliidae | Simulium | *Simulium* spp. |
| Diptera | Empididae | | Empididae spp. |
| Coleoptera | Dytiscidae | | Dytiscidae spp. |
| Coleoptera | Elmidae | Heterlimnius | *Heterlimnius hasegawai* |

**Chubu**

(Fish)

| | | | |
|---|---|---|---|
| Salmoniformes | Salmonidae | Salvelinus | *Salvelinus leucomaenis* spp. |
| Salmoniformes | Salmonidae | Oncorhynchus | *Oncorhynchus masou ishikawai* |

| | | | |
|---|---|---|---|
| Salmoniformes | Salmonidae | Oncorhynchus | *Oncorhynchus mykiss* |
| Perciformes | Cottidae | Cottus | *Cottus pollux* |
| Cypriniformes | Cyprinidae | Pseudogobio | *Pseudogobio* spp. |
| Cypriniformes | Cyprinidae | Opsariichthys | *Opsariichthys platypus* |
| Cypriniformes | Cyprinidae | Candidia | *Candidia temminckii* |
| Cypriniformes | Cyprinidae | Phoxinus | *Phoxinus oxycephalus jouyi* |
| Cypriniformes | Cyprinidae | Phoxinus | *Phoxinus lagowskii steindachneri* |
| Cypriniformes | Cyprinidae | Pseudaspius | *Pseudaspius hakonensis* |
| Cypriniformes | Cobitidae | Cobitis | *Cobitis* spp. |
| Cypriniformes | Cobitidae | Niwaella | *Niwaella delicata* |
| Siluriformes | Amblycipitidae | Liobagrus | *Liobagrus reini* |
| Anguilliformes | Anguillidae | Anguilla | *Anguilla japonica* |
| (Macroinvertebrates) | | | |
| Ephemeroptera | Potamanthidae | Potamanthus | *Potamanthus formosus* |
| Ephemeroptera | Baetidae | Acentrella | *Acentrella gnom* |
| Ephemeroptera | Baetidae | Alainites | *Alainites yoshinensis* |
| Ephemeroptera | Baetidae | Baetiella | *Baetiella japonica* |
| Ephemeroptera | Baetidae | Baetis | *Baetis sahoensis* |

| | | | |
|---|---|---|---|
| Ephemeroptera | Baetidae | Baetis | *Baetis* spp. *J* |
| Ephemeroptera | Baetidae | Baetis | *Baetis taiwanensis* |
| Ephemeroptera | Baetidae | Baetis | *Baetis thermicus* |
| Ephemeroptera | Baetidae | Tenuibaetis | *Tenuibaetis flexifemora* |
| Ephemeroptera | Baetidae | Tenuibaetis | *Tenuibaetis parvipterus* |
| Ephemeroptera | Isonychiidae | Isonychia | *Isonychia valida* |
| Ephemeroptera | Leptophlebiidae | Paraleptophlebia | *Paraleptophlebia* spp. |
| Ephemeroptera | Ameletidae | Ameletus | *Ameletus* spp. |
| Ephemeroptera | Heptageniidae | Cinygmula | *Cinygmula* spp. |
| Ephemeroptera | Heptageniidae | Ecdyonurus | *Ecdyonurus* spp. |
| Ephemeroptera | Heptageniidae | Epeorus | *Epeorus aesculus* |
| Ephemeroptera | Heptageniidae | Epeorus | *Epeorus curvatulus* |
| Ephemeroptera | Heptageniidae | Epeorus | *Epeorus ikanonis* |
| Ephemeroptera | Heptageniidae | Epeorus | *Epeorus latifolium* |
| Ephemeroptera | Heptageniidae | Heptagenia | *Heptagenia* spp. |
| Ephemeroptera | Heptageniidae | Rhithrogena | *Rhithrogena tetrapunctigera* |
| Ephemeroptera | Ephemerellidae | Cincticostella | *Cincticostella orientalis* |
| Ephemeroptera | Ephemerellidae | Cincticostella | *Cincticostella* spp. |
| Ephemeroptera | Ephemerellidae | Drunella | *Drunella* spp. |

| | | | |
|---|---|---|---|
| Ephemeroptera | Ephemerellidae | Ephemerella | *Ephemerella* spp. |
| Ephemeroptera | Ephemerellidae | Teleganopsis | *Teleganopsis punctisetae* |
| Ephemeroptera | Ephemerellidae | Torleya | *Torleya japonica* |
| Ephemeroptera | Ephemeridae | Ephemera | *Ephemera* spp. |
| Plecoptera | Perlodidae | | Perlodidae spp. |
| Plecoptera | Nemouridae | Amphinemura | *Amphinemura* spp. |
| Plecoptera | Nemouridae | Nemoura | *Nemoura* spp. |
| Plecoptera | Nemouridae | Protonemura | *Protonemura* spp. |
| Plecoptera | Perlidae | | Perlidae spp. |
| Plecoptera | Capniidae | | Capniidae spp. |
| Plecoptera | Taeniopterygidae | | Taeniopterygidae spp. |
| Plecoptera | Peltoperlidae | Yoraperla | *Yoraperla uenoi* |
| Plecoptera | Leuctridae | | Leuctridae spp. |
| Plecoptera | Chloroperlidae | | Chloroperlidae spp. |
| Trichoptera | Polycentropodidae | Plectrocnemia | *Plectrocnemia* spp. |
| Trichoptera | Brachycentridae | Micrasema | *Micrasema* spp. |
| Trichoptera | Lepidostomatidae | Lepidostoma | *Lepidostoma* spp. |
| Trichoptera | Philopotamidae | Dolophilodes | *Dolophilodes* spp. |
| Trichoptera | Hydrobiosidae | Apsilochorema | *Apsilochorema sutshanum* |

| | | | |
|---|---|---|---|
| Trichoptera | Psychomyiidae | Psychomyia | *Psychomyia* spp. |
| Trichoptera | Uenoidae | Uenoa | *Uenoa tokunagai* |
| Trichoptera | Sericostomatidae | Gumaga | *Gumaga orientalis* |
| Trichoptera | Apataniidae | Apatania | *Apatania* spp. |
| Trichoptera | Hydropsychidae | Arctopsyche | *Arctopsyche* spp. |
| Trichoptera | Hydropsychidae | Cheumatopsyche | *Cheumatopsyche* spp. |
| Trichoptera | Hydropsychidae | Diplectrona | *Diplectrona* spp. |
| Trichoptera | Hydropsychidae | Hydropsyche | *Hydropsyche albicephala* |
| Trichoptera | Hydropsychidae | Hydropsyche | *Hydropsyche gifuana* |
| Trichoptera | Hydropsychidae | Hydropsyche | *Hydropsyche orientalis* |
| Trichoptera | Hydropsychidae | Hydropsyche | *Hydropsyche setensis* |
| Trichoptera | Hydropsychidae | Macrostemum | *Macrostemum radiatum* |
| Trichoptera | Hydropsychidae | Parapsyche | *Parapsyche* spp. |
| Trichoptera | Rhyacophilidae | Rhyacophila | *Rhyacophila* spp. |
| Trichoptera | Goeridae | Goera | *Goera japonica* |
| Trichoptera | Stenopsychidae | Stenopsyche | *Stenopsyche marmorata* |
| Trichoptera | Stenopsychidae | Stenopsyche | *Stenopsyche sauteri* |
| Trichoptera | Leptoceridae | Ceraclea | *Ceraclea* spp. |
| Trichoptera | Hydroptilidae | Hydroptila | *Hydroptila* spp. |

| | | | |
|---|---|---|---|
| Trichoptera | Glossosomatidae | Agapetus | *Agapetus* spp. |
| Trichoptera | Glossosomatidae | Glossosoma | *Glossosoma* spp. |
| Diptera | Blephariceridae | Agathon | *Agathon bilobatoides* |
| Diptera | Blephariceridae | | Blephariceridae spp. |
| Diptera | Empididae | | Empididae spp. |
| Diptera | Pediciindae | Dicranota | *Dicranota* spp. |
| Diptera | Tipulidae | Tipula | *Tipula* spp. |
| Diptera | Psychodidae | Pericoma | *Pericoma* spp. |
| Diptera | Athericidae | Asuragina | *Asuragina caerulescens* |
| Diptera | Athericidae | Atherix | *Atherix ibis japonica* |
| Diptera | Ceratopogonidae | | Ceratopogonidae spp. |
| Diptera | Limoniidae | Antocha | *Antocha* spp. |
| Diptera | Limoniidae | Hexatoma | *Hexatoma* spp. |
| Diptera | Limoniidae | Molophilus | *Molophilus* spp. |
| Diptera | Simuliidae | Prosimulium | *Prosimulium* spp. |
| Diptera | Simuliidae | Simulium | *Simulium* spp. |
| Diptera | Chironomidae | Chironominae | *Chironominae* spp. |
| Diptera | Chironomidae | Diamesinae | Diamesinae spp. |
| Diptera | Chironomidae | Orthocladiinae | Orthocladiinae spp. |

| | | | |
|---|---|---|---|
| Diptera | Chironomidae | Tanypodinae | Tanypodinae spp. |
| Coleoptera | Elmidae | Elminae | *Elminae* spp. |
| Coleoptera | Elmidae | Grouvellinus | *Grouvellinus* spp. |
| Coleoptera | Elmidae | Optioservus | *Optioservus* spp. |
| Coleoptera | Elmidae | Zaitzevia | *Zaitzevia* spp. |
| Coleoptera | Psephenidae | Ectopria | *Ectopria opaca opaca* |
| Coleoptera | Psephenidae | Eubrianax | *Eubrianax granicollis* |
| Coleoptera | Psephenidae | Mataeopsephus | *Mataeopsephus japonicus* |
| Coleoptera | Scirtidae | | Scirtidae spp. |

**# Calculation of the mean cooling effect associated with watershed geology**

In Table S4, Precipi, Air, Agri Elev indicates the mean total summer precipitation, the mean summer air temperature, the mean site elevation, and the mean proportion of agricultural land use in the watershed, respectively. For each study region, we estimated the mean summer water temperature in two examples, including volcanic streams (100 % of the volcanic rocks in the watershed) and non-volcanic streams (0 % of the volcanic rocks in the watershed) by using the mean value of each covariate. The mean cooling effect in each region was derived from the difference in the estimated mean summer water temperature between the two geological types.

**Table S4** Mean cooling effect associated with watershed geology

| Region | Precipi (mm) | Air (℃) | Agri (%) | Elev (m) | Estimated summer WT in volcanic streams (℃) | Estimated summer WT in non-volcanic streams (℃) | Cooling effect (℃) |
|---|---|---|---|---|---|---|---|
| Okhotsk | 151.7 | 18.1 | 0.0 | 453.3 | 16.34 | 13.04 | 3.30 |
| Sorachi | 280.0 | 18.6 | 0.3 | 333.4 | 16.81 | 13.98 | 2.83 |
| Oshima | 334.4 | 20.3 | 0.0 | 110.5 | 17.99 | 15.65 | 2.34 |
| Kantou | 440.7 | 21.1 | 0.8 | 614.3 | 17.61 | 15.74 | 1.87 |
| Chubu | 990.8 | 21.4 | 0.9 | 483.3 | 16.77 | 16.52 | 0.25 |

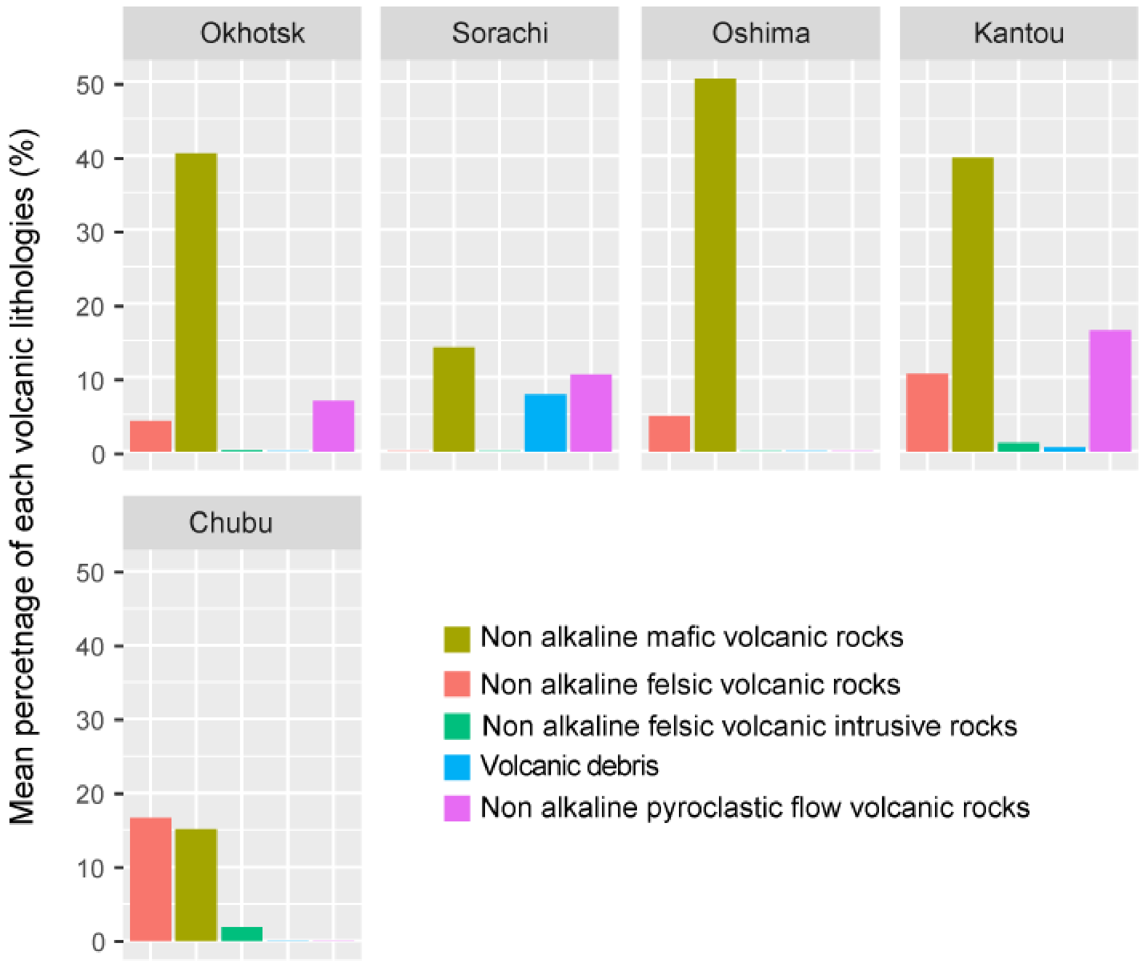


**Fig. S1** Mean proportion of every volcanic lithology in the volcanic geology of each study region.

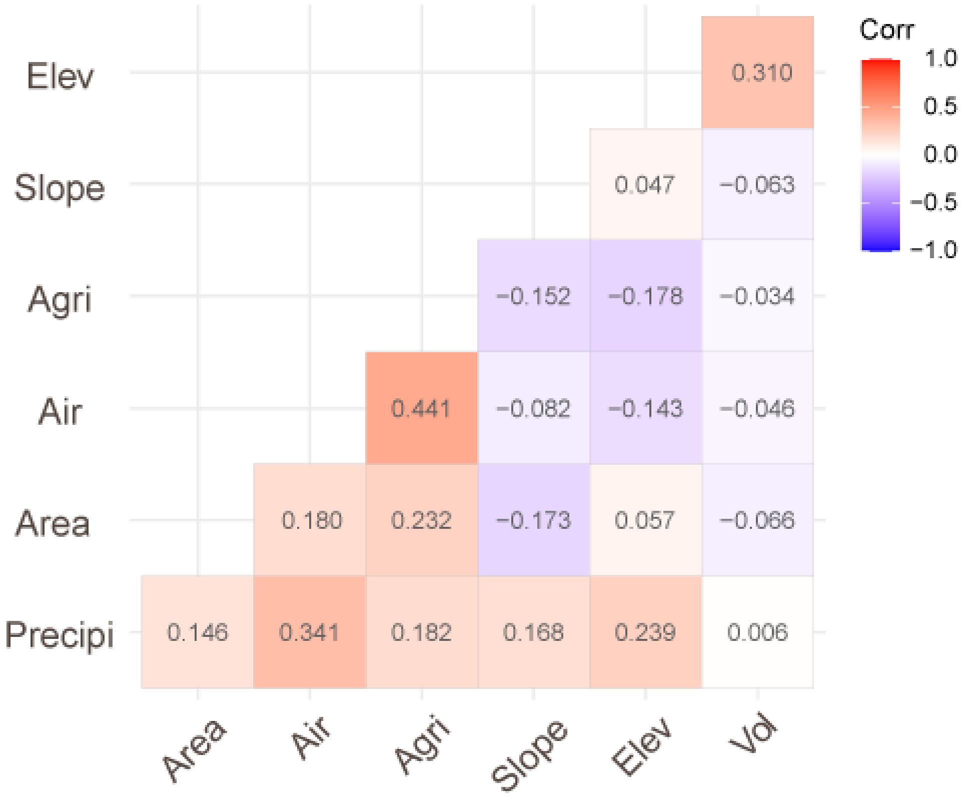


**Fig. S2** Correlations among covariates for the water temperature modeling. Elev: site elevation (m), Slope: site slope, Agri: the proportion of agricultural land use in the watershed (%), Air: mean summer air temperature (°C), Area: Drainage area ($km^2$), Precipi: Total summer precipitation (mm) and Vol: Volcanic rocks in the watershed (%).

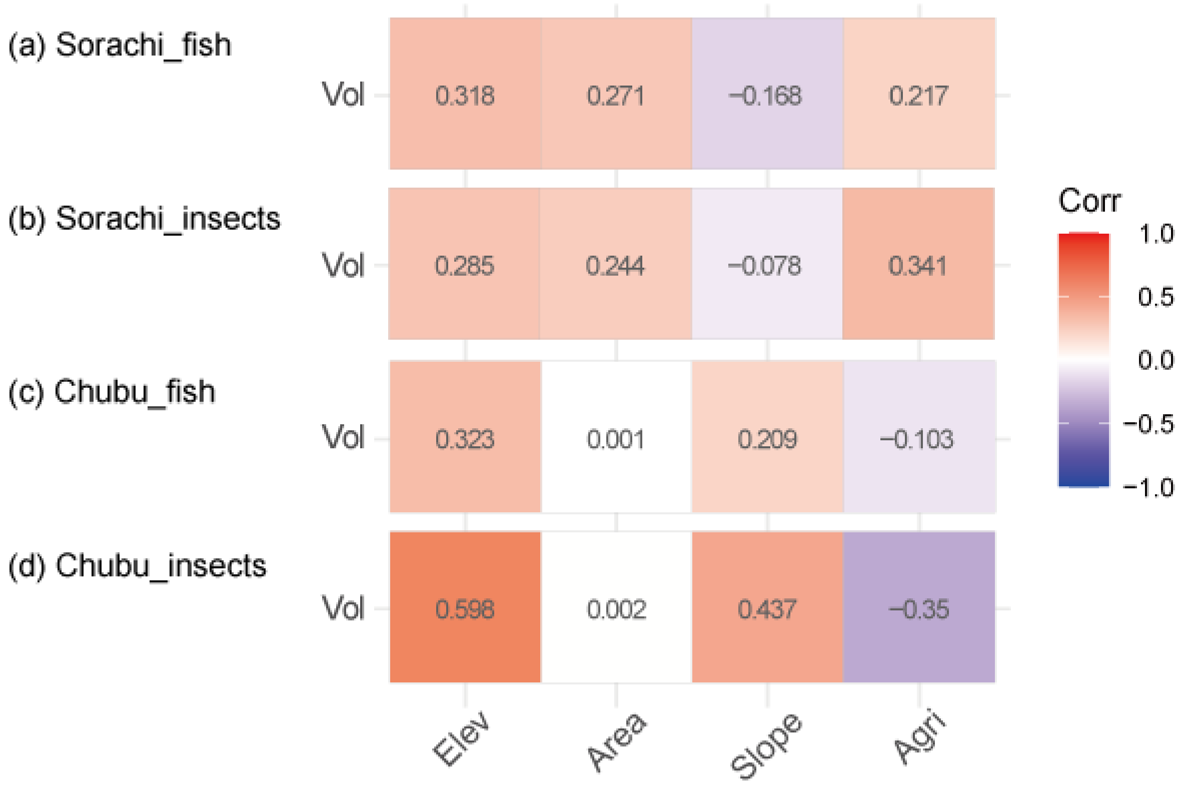


**Fig. S3.** Correlations of watershed geology with other major landscape-scale factors. (a) Fish monitoring sites in the Sorachi, (b) macroinvertebrates monitoring sites in the Sorachi, (c) fish monitoring sites in the Chubu and (d) macroinvertebrates monitoring sites in the Chubu. Vol: Volcanic rocks in the watershed (%), Elev: Site elevation (m), Area: Drainage area ($km^2$), Slope: site slope, and Agri: the proportion of agricultural land use in the watershed (%).

### Analyses for the occurrence of *C. pollux*

Sampling was conducted from 2015 to 2017 in 32 sites in 32 streams, which are in the Nagara, Ibi and Kiso rivers of the Chubu region. We selected low order (area of watershed < 100 $km^2$) and low disturbance (forest cover ratio in the watershed > 80%) streams as the study sites. Following Suzuki et al. (2021) testing the relationship between the occurrence of *C. nozawae* and the summer water temperature, we analyzed the occurrence of *C. pollux* using GLMs with a binomial error and logit link function. The response variable was the presence-absence of *C. pollux*. The explanatory variables were the mean summer water temperature, stream slope, catchment size, and the log-transformed proportion of farmland in the catchments. We also incorporated the survey area of each sampling site in the models to account for variability in the sampling efforts. We constructed models for all cases with a best-subset procedure, and the model performance was evaluated based on the AICc. We conducted model averaging using superior models (ΔAICc < 2) and applied the parameters for which the 95% CIs did not include 0 to the final model. We found that the mean summer water temperature was the sole influential factor and showed a negative relationship with the occurrence of *C. pollux*. This occurrence pattern was the same as that of *C. nozawae* reported by Suzuki et al. (2021). In the final model, the occurrence probability of 0.5 for *C. pollux* corresponded to 18.4 °C.

Suzuki, K., Ishiyama, N., Koizumi, I., & Nakamura, F. (2021). Combined Effects of Summer Water Temperature and Current Velocity on the Distribution of a Cold-Water-Adapted Sculpin (*Cottus nozawae*). Water, 13(7), 975.

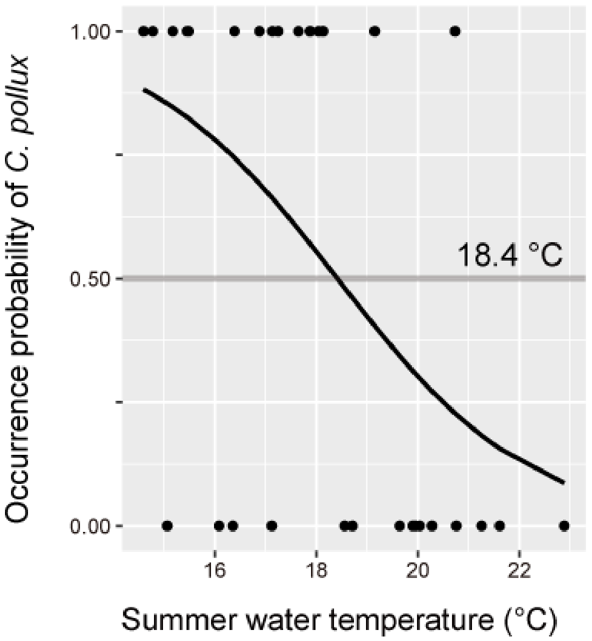


**Fig. S4** The relationship between the mean summer water temperature and the probability of *C. pollux* occurrence. The black circles and line indicate the observed data and the predicted probability, respectively. The final model yielded an $R^2$ of 0.23 and an area under the receiver operating characteristic curve (AUC) of 0.77.

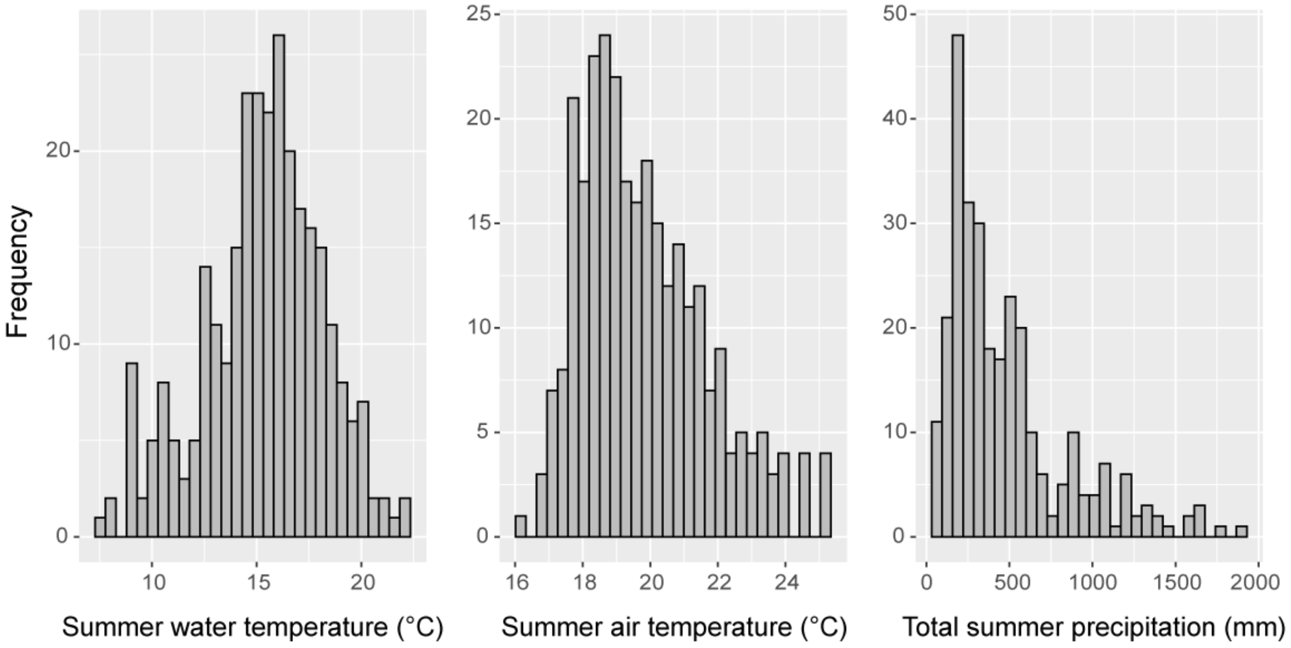


**Fig. S5** Frequency of the observed mean summer water temperatures and climatic conditions at each monitoring site.

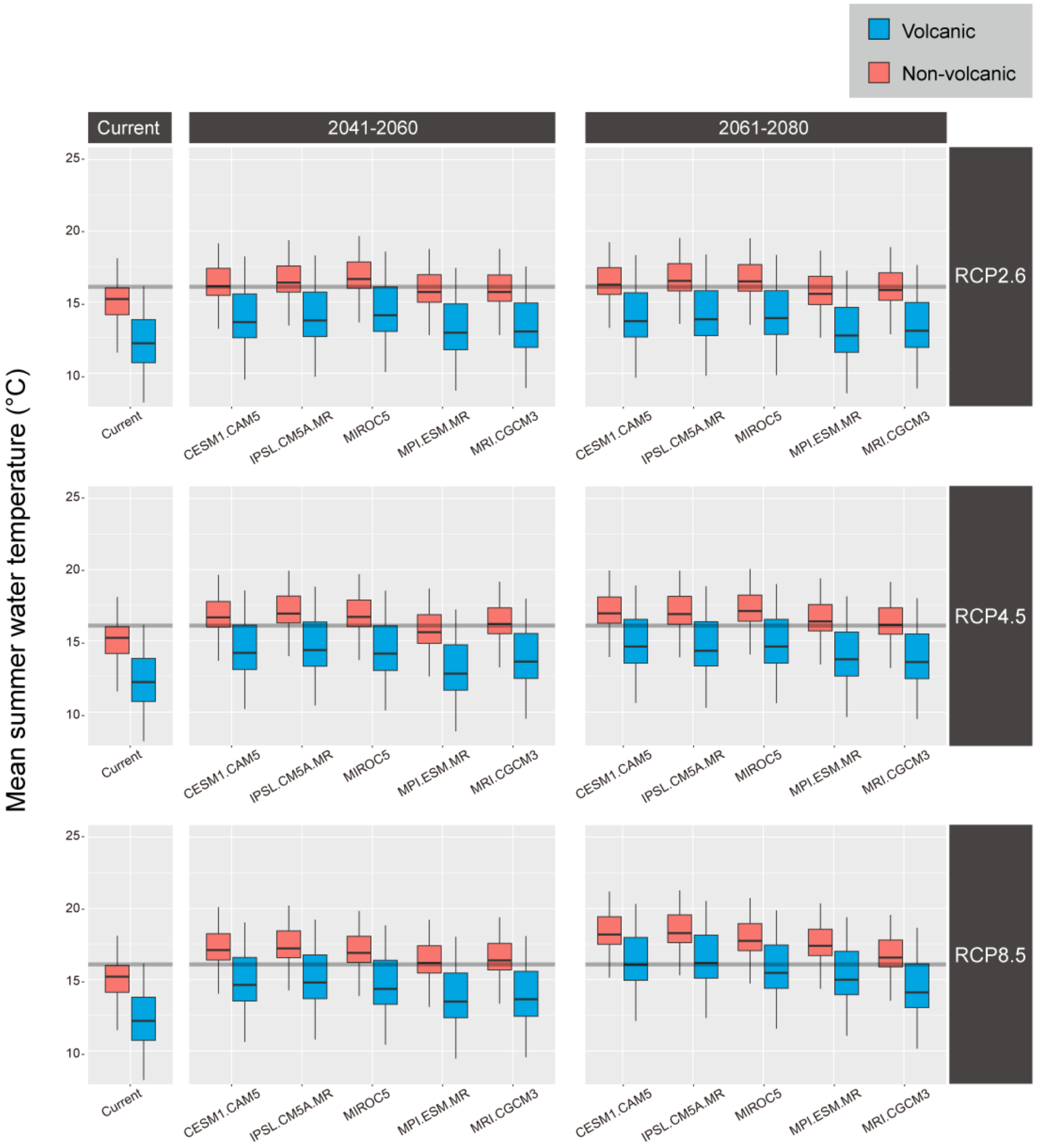


**Fig. S6** Current and future mean summer water temperatures in northern Japan (Okhotsk, Sorachi, Oshima), and their relation to the suitable thermal limit for *C. nozawae.* Predictions are shown by each GCMs, period and RCP. The lines in the middle of boxes represent the median, and the lower and upper ends of boxes are the 25% and 75% quartiles respectively. The thick gray line is the limit of the preferable temperature (16.1 °C).

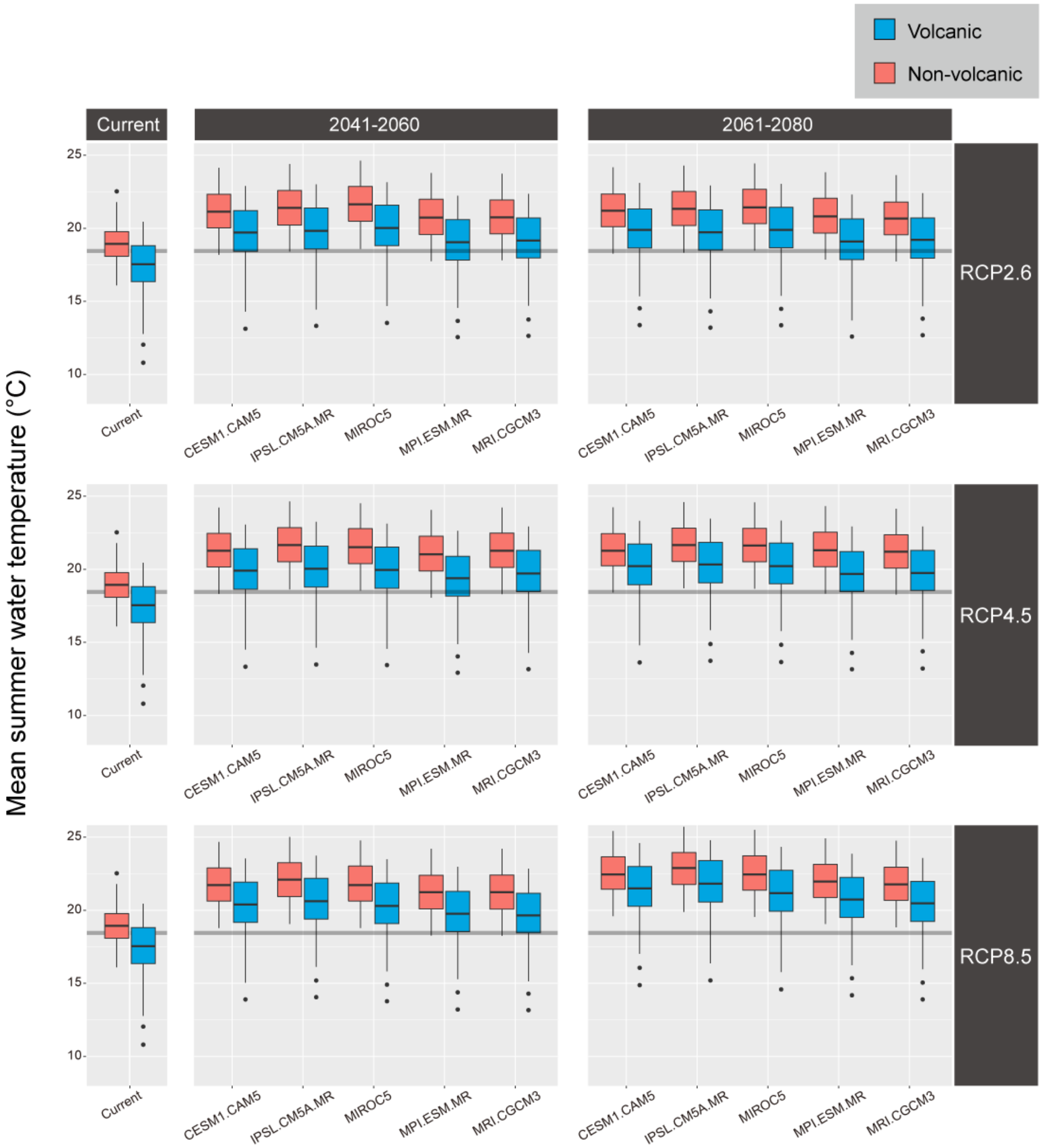


**Fig. S7** Current and future mean summer water temperatures in central Japan (Kantou, Chubu), and their relation to the suitable thermal limit for *C. pollux*. Predictions are shown by each GCMs, period and RCP. The lines in the middle of boxes represent the median, and the lower and upper ends of the boxes are the 25% and 75% quartiles respectively. The thick gray line is the suitable thermal limit (18.4 °C).

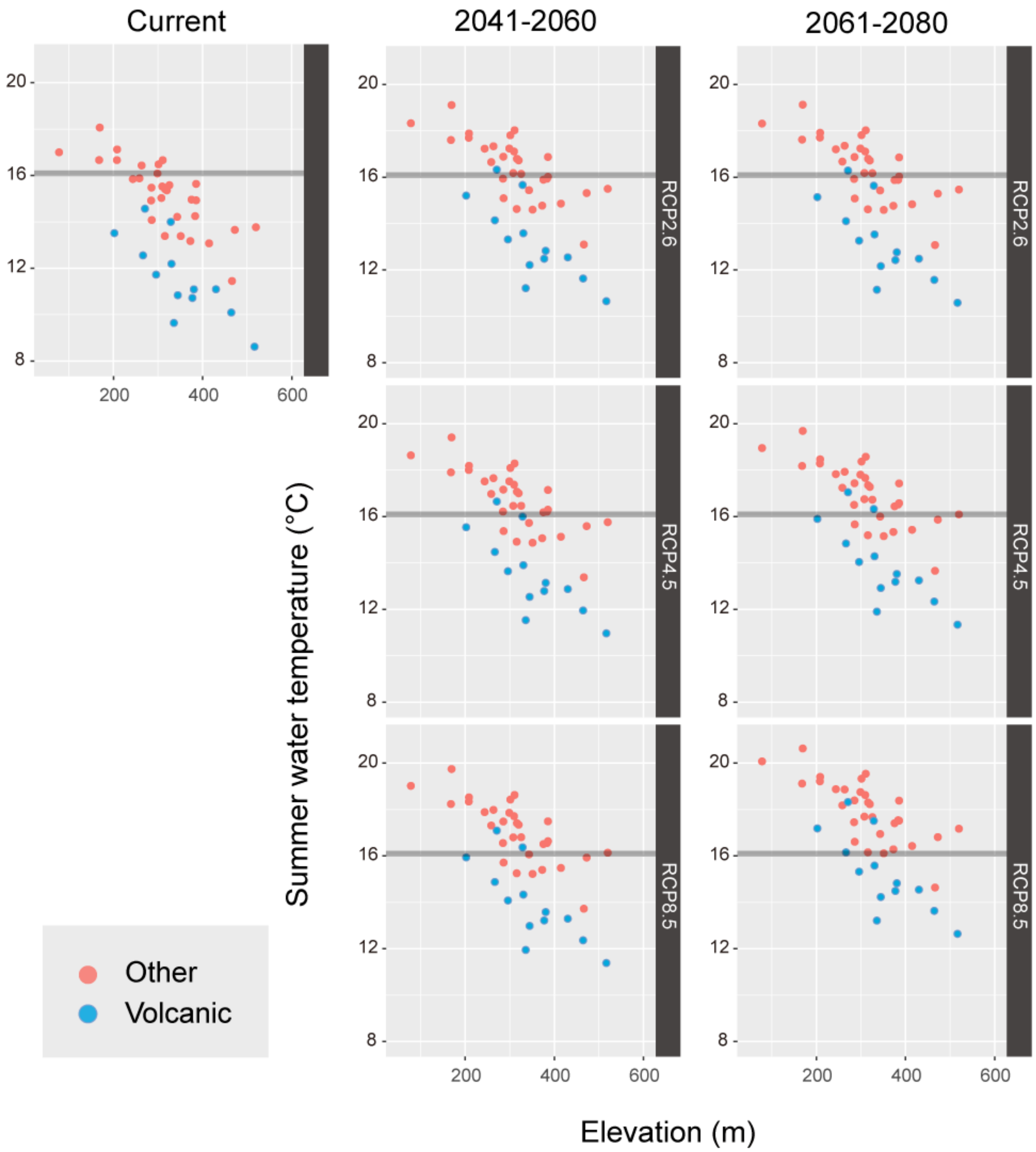


**Fig. S8** The relationship between the site elevation and the summer water temperatures in the Sorachi river under contemporary and future climates. The thick gray line is the thermal threshold for *C. nozawae* (16.1 °C). The relationships are shown by each period and the RCP.
